# Broadly neutralizing antibodies to SARS-related viruses can be readily induced in rhesus macaques

**DOI:** 10.1101/2021.07.05.451222

**Authors:** Wan-ting He, Meng Yuan, Sean Callaghan, Rami Musharrafieh, Ge Song, Murillo Silva, Nathan Beutler, Wilma Lee, Peter Yong, Jonathan Torres, Mariane Melo, Panpan Zhou, Fangzhu Zhao, Xueyong Zhu, Linghang Peng, Deli Huang, Fabio Anzanello, James Ricketts, Mara Parren, Elijah Garcia, Melissa Ferguson, William Rinaldi, Stephen A. Rawlings, David Nemazee, Davey M. Smith, Bryan Briney, Yana Safonova, Thomas F. Rogers, Shane Crotty, Darrell J. Irvine, Andrew B. Ward, Ian A. Wilson, Dennis R. Burton, Raiees Andrabi

**Author notes:** Corresponding author. (I.A.W); (D.R.B.); (R.A.). These authors contributed equally to this work (Wan-ting He, Meng Yuan, Sean Callaghan).

## Abstract

To prepare for future coronavirus (CoV) pandemics, it is desirable to generate vaccines capable of eliciting neutralizing antibody responses against multiple CoVs. Because of the phylogenetic similarity to humans, rhesus macaques are an animal model of choice for many virus-challenge and vaccine-evaluation studies, including SARS-CoV-2. Here, we show that immunization of macaques with SARS-CoV-2 spike (S) protein generates potent receptor binding domain cross-neutralizing antibody (nAb) responses to both SARS-CoV-2 and SARS-CoV-1, in contrast to human infection or vaccination where responses are typically SARS-CoV-2-specific. Furthermore, the macaque nAbs are equally effective against SARS-CoV-2 variants of concern. Structural studies show that different immunodominant sites are targeted by the two primate species. Human antibodies generally target epitopes strongly overlapping the ACE2 receptor binding site (RBS), whereas the macaque antibodies recognize a relatively conserved region proximal to the RBS that represents another potential pan-SARS-related virus site rarely targeted by human antibodies. B cell repertoire differences between the two primates appear to significantly influence the vaccine response and suggest care in the use of rhesus macaques in evaluation of vaccines to SARS-related viruses intended for human use.

**ONE SENTENCE SUMMARY:** Broadly neutralizing antibodies to an unappreciated site of conservation in the RBD in SARS-related viruses can be readily induced in rhesus macaques because of distinct properties of the naïve macaque B cell repertoire that suggest prudence in the use of the macaque model in SARS vaccine evaluation and design.

## MAIN

The rapid development of multiple vaccines to counter the COVID-19 pandemic caused by SARS-CoV-2 has been a triumph for science and medicine. The reason for these successes is likely due, in part, to the relative ease of inducing protective neutralizing antibodies (nAbs) in humans by immunization with the spike (S) protein, the key component of most vaccines (*1–4*). Different presentations of S in the vaccines induce relatively high serum nAb responses in most individuals that appear to strongly target the ACE2 receptor binding site (RBS) in the receptor binding domain (RBD) (*1, 5–9*). Natural infection with SARS-CoV-2 targets the RBD similarly (*1, 10–22*). A key contributor is the human VH3-53 antibody germline gene that encodes a major subset (RBS-A, Class 1) of highly potent immunodominant nAbs that require minimal somatic hypermutation (SHM) and are specific for SARS-CoV-2 (*22–25*). These and other nAbs to the RBS are likely responsible for driving some of the mutations found in Variants of Concern (VOCs) (*23, 26–30*). Here, we immunized rhesus macaques with SARS-CoV-2 S-protein to compare nAb responses between macaques and humans with a particular emphasis on the breadth of elicited neutralizing responses to sarbecoviruses and the effects of differences between the naive antibody repertoires of the two primate species on responses.

### SARS-CoV-2 S-protein prime-boost immunization leads to robust sarbecovirus broadly neutralizing antibody (bnAb) responses in rhesus macaques

A recombinant prefusion-stabilized soluble S-protein in a saponin adjuvant (SMNP) was administered through subcutaneous injection (s.c.) in naïve rhesus macaques (RMs) (n = 8). Since the manner of immunogen administration may shape immune responses (*31–33*), two protocols were evaluated: a conventional prime-boost consisting of bolus injections (100µg) at weeks 0 and 10 for Group 1 animals (n = 4), and an escalating dose (ED) prime and bolus boost for Group 2 (n=4) (Fig. 1A). Serum Ab responses to SARS-CoV-2 S-protein, evaluated by enzyme-linked immunosorbent assay (ELISA) (figs. S1 and S2), were detected in both groups at week 2 and, although responses at week 4 were stronger for the ED group, no major differences were found subsequently. Boost immunization at week 10 increased EC_50_ binding titers to the 10^3^-10^4^ range for all animals, indicating a strong antibody recall response. ID_50_ neutralization titers showed a similar pattern where both immunization groups developed specific neutralizing antibody responses by week 4, which were enhanced by week 12, post-boost (figs. S1 and S2).

**Fig. 1.**
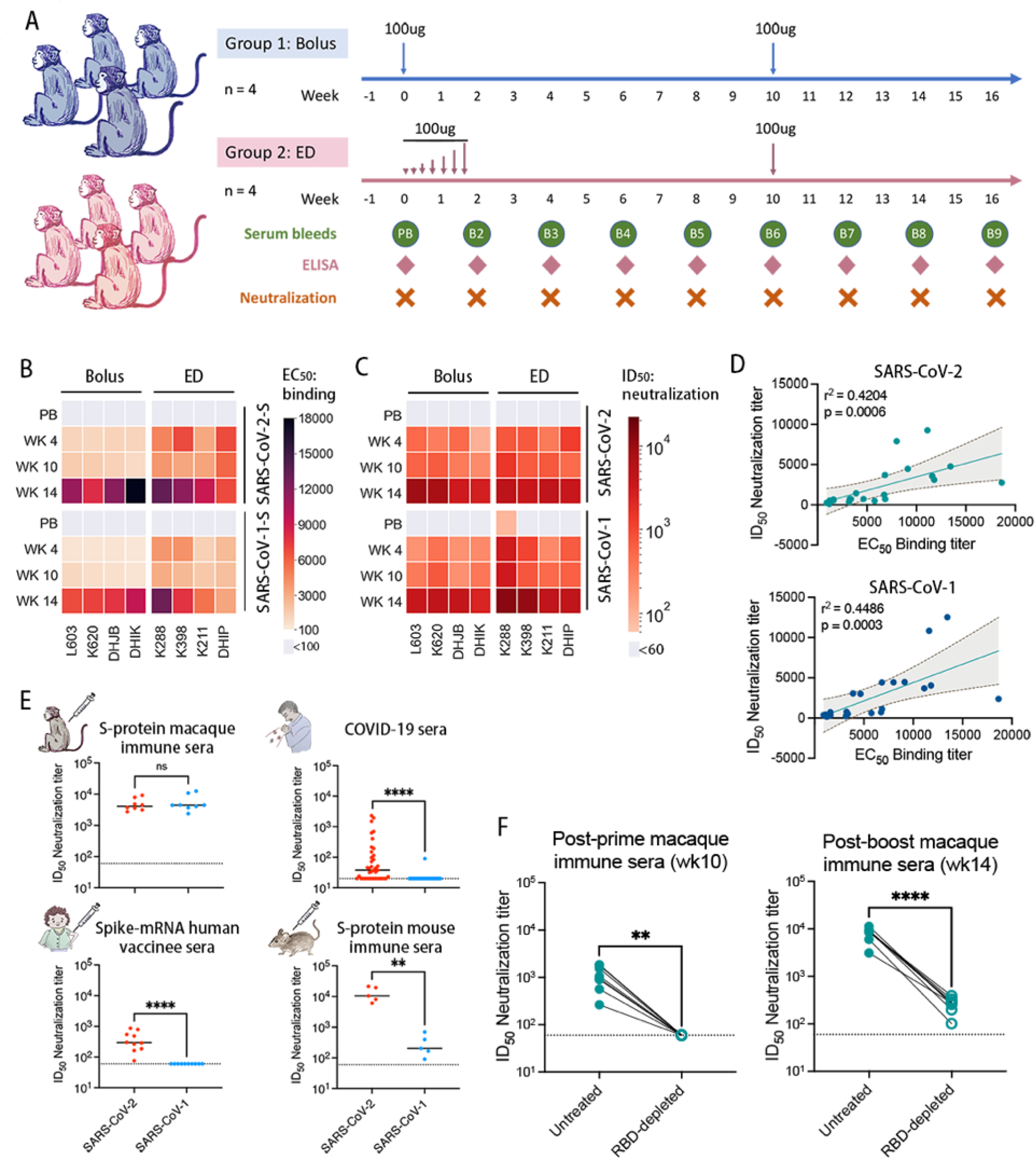
SARS-CoV-2 and SARS-CoV-1 cross-reactive binding and neutralizing antibody responses in SARS-CoV-2 S protein-immunized rhesus macaques. **(A)** SARS-CoV-2 S protein prime-boost immunization in rhesus macaques and sampling schedule. Animals were primed (wk0) with 100µg of SARS-CoV-2 S protein along with adjuvant SMNP, by bolus (Group 1: n = 4) and escalating dose (ED) (Group 2: n = 4) immunization and both groups further boosted (wk10) with 100µg of the S protein immunogen. Serum and peripheral blood mononuclear cells (PBMCs) were collected every two weeks for immune response analysis. **(B)** Heatmap showing EC_50_ ELISA binding of SARS-CoV-2 S protein-immunized NHP sera from bolus and dose escalation schemes against different S proteins. EC_50_ binding responses for pre-bleed (PB), post- prime (wk4 and wk10) and post-boost (wk14) sample time points are shown. The SARS-CoV-2 S protein-immunized sera exhibit a strong cross-reactivity with SARS-CoV-2 and SARS-CoV-1 S- proteins. **(C)** SARS-CoV-2 and SARS-CoV-1 specific serum ID_50_ neutralizing antibody titers in SARS-CoV-2 S protein-immunized rhesus macaques. ID_50_ neutralizing antibody responses are shown for pre-bleed (PB), post prime (wk4 and wk10), and post boost (wk14) sample time points. **(D)** NHP immune sera binding to SARS-CoV-2 or SARS-CoV-1 S proteins correlated modestly with neutralization against the corresponding virus. SARS-CoV-2 or SARS-CoV-1 EC_50_ serum antibody binding titers and ID_50_ neutralizing antibody titers were compared by nonparametric Spearman correlation two-tailed test with 95% confidence interval. The Spearman correlation coefficient r and the p-value are indicated. **(E)** Comparison of cross-neutralizing activities of SARS-CoV-2 S protein-immunized NHP immune sera (wk14, after two immunizations at wk0 and wk10) and COVID-19 sera from SARS-CoV-2-infected individuals, sera from mRNA S protein- immunized individuals and S-protein immunized mice. Statistical comparisons between groups were performed using a Mann-Whitney two-tailed test, (**p <0.01; ****p < 0.0001; ns- p >0.05). **F.** Neutralization of SARS-CoV-2 by post-prime (wk10) and post-boost (wk14) macaque immune sera and the corresponding anti-RBD antibody-depleted sera (RBD-depleted). Anti-RBD polyclonal antibodies in the immune sera were depleted by adsorption on recombinant monomeric RBD and the depleted sera were tested for SARS-CoV-2 neutralization. The SARS-CoV-2 ID_50_ neutralizing antibody titers for untreated and RBD-depleted sera were compared by Mann- Whitney two-tailed test, (**p <0.01; ****p < 0.0001).

Next, we examined serum responses for cross-reactivity with SARS-CoV-1 (Fig. 1B, fig. S2). Strong cross-reactive binding responses to SARS-CoV-1 S protein were observed (Fig. 1B, fig. S2), as well as strong cross-neutralizing antibody responses against SARS-CoV-1 (Fig. 1C, fig. S2). ID_50_ neutralization titers showed a modest correlation with EC_50_ binding titers for both SARS-CoV-2 and SARS-CoV-1 (Fig. 1D). The elicitation of potent cross-neutralizing responses in rhesus macaques by S-protein immunization is in stark contrast to human SARS-CoV-2 natural infection, which typically results in autologous nAb responses where cross-neutralizing activity against SARS-CoV-1 is rare (Fig. 1E, fig. S3) (*34–36*). In humans vaccinated with mRNA S-protein, we similarly found SARS-CoV-2 autologous but not SARS-CoV-1 cross-reactive nAb responses (Fig. 1E, fig. S3) as also described by others (*37*).

To examine the role of species-specific B cell immunogenetic differences, we vaccinated a group of 5 mice twice (wk0-prime, wk3-boost) with SARS-CoV-2 S-protein and SMNP adjuvant (fig. S4). Consistent with earlier studies (*38–40*), SARS-CoV-2 S protein immunization of mice elicited high titers of autologous SARS-CoV-2 nAbs; however, the SARS-CoV-1 cross-neutralizing antibody titers were significantly less than the macaque cross-nAb responses (Fig. 1E, fig. S4), indicating a strong species-dependent contribution to development of bnAbs to SARS-CoV-2 S protein.

### Specificities and breadth of neutralization of the polyclonal macaque nAbs

We next characterized polyclonal nAb specificities generated in the macaques by measuring the binding of immune sera in the presence of three well-characterized human nAbs to SARS-CoV-2 S-protein and RBD by Bio-layer Interferometry (fig. S5). These nAbs define unique epitopes on the RBD and were isolated from a COVID-19 convalescent donor (CC12.3 and CC12.19) (*11, 22*) and a SARS-CoV-1 donor (CR3022) (*11, 41*). CC12.3 is a VH3-53 nAb targeting the ACE2 binding site and designated as RBS-A or class 1 (*22–24*). CC12.19 recognizes a complex RBD epitope but competes with some non-RBD Abs (*11*). CR3022 recognizes the class 4 epitope site (*23, 24*). Macaque sera bound well to S-protein in the presence of the nAbs suggesting the RBD nAb response was a minor part of the total S-response (fig. S5). However, binding to RBD was strongly inhibited at week 10 by all 3 nAbs consistent with a strong response to RBD neutralizing epitopes (fig. S5). The inhibition was reduced following boosting suggesting that antibody responses to B cell epitopes outside of the RBD may increase after secondary immunization (*42, 43*).

To further determine the contributions of RBD-specific antibodies in polyclonal immune sera, we used recombinant RBD to deplete sera of RBD antibodies and assessed neutralization at week 10 (post-prime) and week 14 (post-boost). Depletion eliminated neutralization of week-10 sera and greatly reduced activity of week-14 sera (fig. 1F). Depletion of anti-RBD antibodies was confirmed by BLI binding studies with SARS-CoV-2 RBD and S-protein (fig. S6). The results suggest that SARS-CoV-2 neutralization in both post-prime and post-boost macaque immune sera is dominated by RBD-specific nAb responses, but post-boost nAbs to non-RBD S epitopes or trimer-dependent RBD epitopes may minimally contribute to neutralization.

Since macaque sera competed strongly with human RBD nAbs and neutralization was largely RBD directed, we investigated other sarbecoviruses by engrafting the RBD from those viruses onto the SARS-CoV-2 backbone (*44–46*). We chose the 3 sarbecoviruses, WIV1, RaTG13 and pang17, which use ACE2 receptor for cell entry and are phylogenetically related to SARS-CoV-2 and SARS-CoV-1 (Fig. 2A, fig. S7). The week-14 macaque immune sera consistently and potently neutralized all three chimeric pseudoviruses (Fig. 2B) with the WIV1 chimera being most effectively neutralized averaged over all animals. COVID-19 sera only neutralized the closely related RaTG13 chimera at a lower level than the macaque sera (Fig. 2B). Spike mRNA human immune sera showed moderate neutralization against WIV1 and RaTG13 chimeras and sporadic neutralization of the pang17 chimera. The results suggest the macaque immune sera targets more conserved RBD regions on sarbecoviruses than sera from human infection or human mRNA vaccination. The fine details of this cross-reactivity are consistent with the relatedness of these viruses to SARS-CoV-1 and SARS-CoV-2. RaTG13 RBD is closer to SARS-CoV-2 (fig S7) and is neutralized more efficiently by COVID-19 sera in which neutralization is predominantly RBD directed. WIV1 is closer to SARS-CoV-1 and is neutralized more potently by the macaque immune sera that neutralizes SARS-CoV-1 more effectively than SARS-CoV-2.

**Fig. 2.**
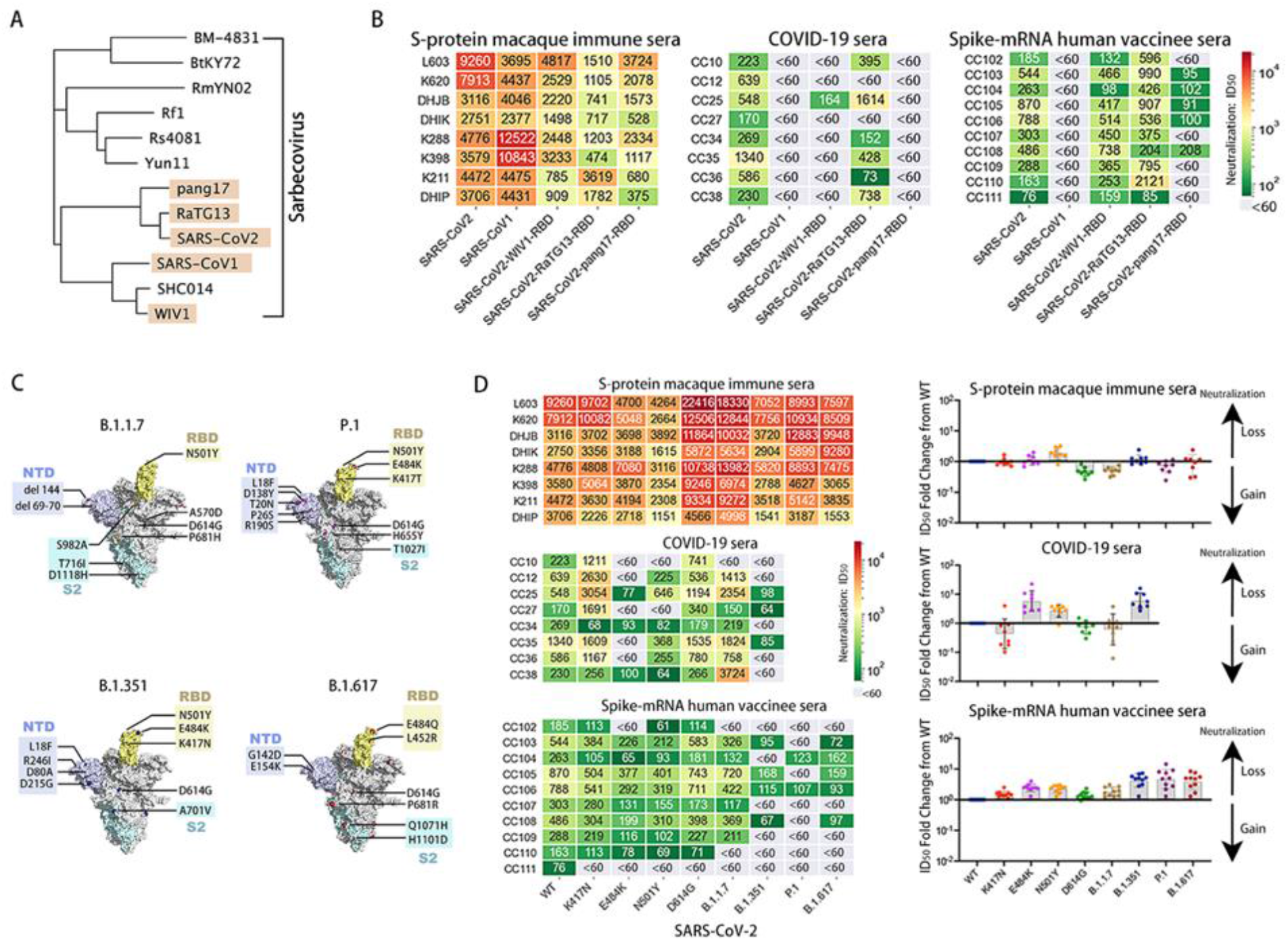
SARS-CoV-2 S-protein immunized macaque sera show neutralization with sarbecovirus spike RBD-swapped chimeric SARS-CoV-2 pseudotyped viruses and SARS-CoV-2 variants circulating in humans. **(A)** Phylogenetic tree of 12 spike glycoproteins representing major sarbecovirus lineages. The five taxa highlighted in beige are SARS-like viruses, related by a common ancestor, that circulate in humans, bats and pangolins. Highlighted strains that produced infectious viruses in an hACE2-expressing reporter assay were selected for the characterization of cross-reactive neutralizing antibody responses. **(B)** Heatmap showing ID_50_ neutralizing antibody titers of SARS-CoV-2 S-protein-immunized wk14 macaque immune sera (left), COVID-19 convalescent sera (middle), and spike-mRNA human vaccinee sera with SARS-CoV-2, SARS-CoV-1, and RBD-swapped WIV1, RaTG13, and pang17 chimeric SARS-CoV-2 pseudoviruses. ID_50_ neutralization titers less than 60 are shown in gray. **(C)** SARS-CoV-2 spike variants of concern (VOCs), B.1.1.7 (alpha), B.1.351 (beta), P.1 (gamma) and B.1.617 (delta), annotated using the cryo-EM structure (PDB:7A94). A single RBD domain in the up configuration is colored in yellow, and the corresponding NTD and S2 domain are colored in light blue and pale cyan, respectively. All mutations and deletions in the spikes of VOCs are labeled accordingly. **(D)** ID_50_ neutralization titers of the antibodies from wk14 SARS-CoV-2 S-protein-immunized macaque immune sera, COVID-19 convalescent sera, and spike-mRNA human vaccinee sera for SARS-CoV-2 incorporating RBD mutations and for variant viruses, B.1.1.7, B.1.351, P.1 and B.1.617. The ID_50_ titers are shown in the heatmap on the left and the fold change of the ID_50_ compared with WT (Wuhan) SARS-CoV-2 are shown in the dot plot on the right. The loss or gain of neutralization potency is indicated by the arrow on the right side of the dot plot.

Next, we compared the macaque, human COVID-19 and mRNA vaccine sera for neutralization of SARS-CoV-2 incorporating key mutations from current SARS-CoV-2 VOCs, B.1.1.7 (alpha), B.1.351 (beta), P.1 (gamma) and B.1.617 (delta) (Fig. 2C) (*9, 23, 28, 29*). While human COVID- 19 and vaccine sera showed significant loss of neutralization activity against point mutants and SARS-CoV-2 VOCs, macaque immune sera maintained neutralizing activity (Fig. 2D). This result is again consistent with macaque sera targeting more conserved regions of the S-protein and apparently thereby yielding antibodies with less sensitivity to known escape mutations.

### Isolation and neutralizing properties of monoclonal antibodies from immunized macaques

We isolated mAbs from two animals (K288 and K398) by sorting antigen-specific single B cells using SARS-CoV-2 and SARS-CoV-1 S proteins as baits (Fig 3A). Briefly, CD19^+^CD20^+^ IgG^+^ IgM^-^ double positive B cells for SARS-CoV-2 and SARS-CoV-1 S-protein were sorted from peripheral blood mononuclear cells (PBMCs) from immunized animals. Paired heavy (HC) and light (LC) chain sequences from single B cells were recovered and 40 mAbs were expressed (Fig. 3B). BLI binding revealed that all but 6 mAbs exhibited binding to SARS-CoV-2 and/or SARS-CoV-1 S-protein (Fig. 3B, fig. S8). 14 of 34 mAbs showed cross-neutralization of SARS-CoV-1 and SARS-CoV-2, while 3 mAbs neutralized only SARS-CoV-2 (Fig. 3B, fig. S8). Unexpectedly, most cross-neutralizing mAbs were more potent against SARS-CoV-1 than SARS-CoV-2 (Fig 3B, fig. S8). 17 of 34 mAbs bound S-protein only and not RBD or NTD, but none showed neutralization. All neutralizing mAbs (except one NTD-specific) bound strongly to RBDs (Fig. 3B, fig. S8). Overall, these results are consistent with the polyclonal nAb data described above (Fig. 1) and show that macaque immunization here with S-protein generates potent SARS-CoV-1/2 nAb responses.

**Fig. 3.**
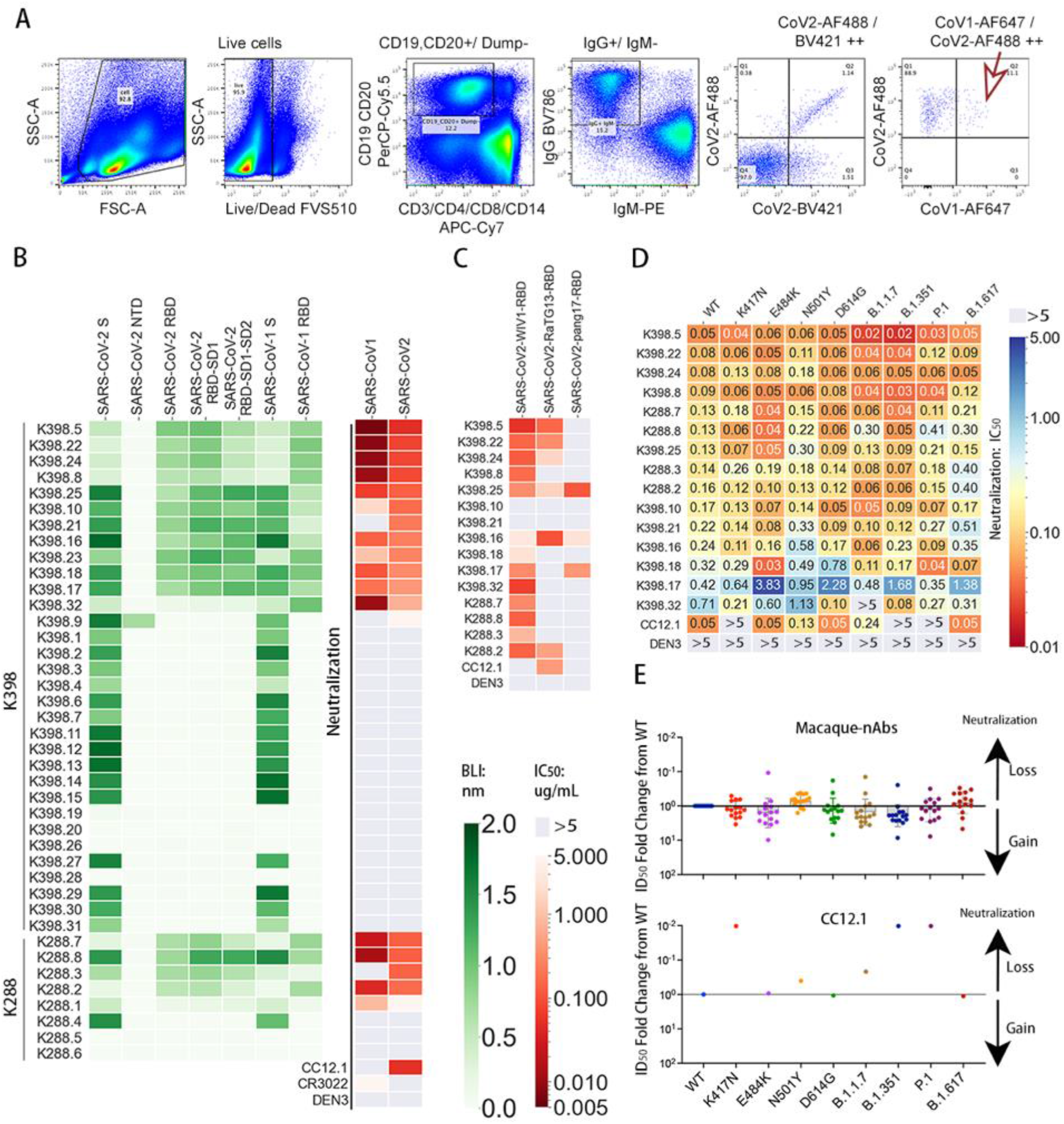
Isolation and characterization of cross-neutralizing monoclonal antibodies from two SARS-CoV-2 S-protein-immunized rhesus macaques. **(A)** Flow cytometry plots showing the sorting strategy used to isolate single isotype-switched B cells that specifically bind to both SARS-CoV-1 and SARS-CoV-2 S-protein fluorescently labelled probes. The class-switched memory B cells were gated as SSL, CD3^−^, CD4^−^, CD8^−^, CD14^−^, CD19^+^/CD20^+^, IgM^−^, IgG^+^. The antigen specific IgG+ B cells selected, as indicated by a red arrow were used to obtain paired heavy and light chain Immunoglobulin sequences for mAb expression and characterization. **(B)** Heatmap showing the BLI binding responses of isolated monoclonal antibodies from rhesus macaques, K398 and K288 against SARS-CoV-2 S-protein, SARS-CoV-2 S-protein-derived domains and subdomains, NTD, RBD, RBD-SD1, RBD-SD1-2, SARS-CoV-1 S-protein and SARS-CoV-1-RBD (indicated with a green color scale). The IC_50_ neutralizing titer against SARS-CoV-1 and SARS-CoV-2 pseudoviruses are shown in the right columns with a red color scale. **(C)** Neutralization of macaque cross-neutralizing mAbs against SARS-CoV-2 pseudoviruses with the RBDs swapped with those of sarbecoviruses identified in bat and pangolin. IC_50_s are color-coded and an IC_50_ > 5µg/mL is indicated in gray. **(D)** IC_50_ of macaque cross-neutralizing mAbs against the pseudoviruses with single amino acid mutations in the SARS-CoV-2 spike as well as the major variants of concern (B.1.1.7, B.1.351, P.1 and B.1.617) circulating in humans. IC_50_ > 5µg/mL are shown in gray. **(E)** Neutralization IC_50_ fold changes of mAbs with mutants compared with the Wuhan SARS-CoV-2 pseudovirus are shown in the dot plots with geometric mean labeled by the bar in gray. The gain or loss of neutralizing potency is indicated by the arrows on the right.

We next extended the neutralization studies to a range of sarbecovirus RBD chimeric SARS-CoV-2 pseudoviruses (Fig. 3C, fig. S8). Most of the mAbs that showed potent SARS-CoV1/2 neutralization also neutralized WIV1 but activity against RaTG13 and pang17 was more sporadic. Some mAbs did neutralize all 5 sarbecoviruses tested, albeit with relatively lower neutralization potency (Fig. 3C, fig. S8). The cross-reactivity of mAbs was also confirmed by BLI binding studies with monomeric RBDs derived from spike of WIV1, RatG13 and Pang17 sarbecoviruses (fig. S9).

We also tested neutralization by these cross-reactive mAbs against SARS-CoV-2 VOCs and key mutants. Consistent with the macaque sera mapping above, the mAbs were generally equally effective against variants (Fig. 3, D and E). In contrast, CC12.1 isolated from a COVID-19 human donor showed complete loss of neutralizing activity with the SARS-CoV-2 K417N variant and B.1.351 and P.1 circulating escape variants (Fig. 3, D and E).

### Properties of mAbs associated with broad neutralization

We next sought to determine molecular differences between mAbs that would explain their differential neutralization. First, we analyzed the mAb sequences and found that they utilized a range of germline gene families with strong enrichment for IGHV3-73 and, to some extent, IGHV3- 50 and IGHV5-15, heavy chain germline genes (Fig. 4A, fig. S10). None of the IGHV3-73 Abs from either animal (K288 and K398) were clonal members, suggesting independent selection of this gene family with common VH-germline gene features (fig. S10). Notably, the rhesus macaque IGHV3-73 germline gene is closely related in sequence to human IGHV3-53 and IGHV3-66 genes, commonly utilized by human SARS-CoV-2 nAbs isolated from natural infection and vaccination, that initially suggested the possibility of convergent modes of RBD site epitope recognition (fig. S11) (*9, 20, 22, 23, 25*). Some cross-nAb members were encoded by non-IGHV3- 73 gene families (Fig. 4, A and B, fig. S10). However, most of the IGHV3-50 and IGHV5-15 encoded antibodies were non-neutralizing, including some clonally expanded antibody lineages isolated from animal K398 (Fig. 4, A and B, fig. S10). The CDRH3 lengths ranged from 9-23 amino acid residues (Fig. 4C, fig. S10) with nAbs possessing relatively longer CDRH3 loops (median = 17aa) than non-nAbs (median = 12aa). Two nAbs used the longest CDRH3 of 23aa residues (fig. S10). The nAbs had relatively lower SHM compared to non-nAbs (median nt SHMs; neutralizing = 10, non-neutralizing = 14.5) (Fig. 4C, fig. S10).

**Fig. 4.**
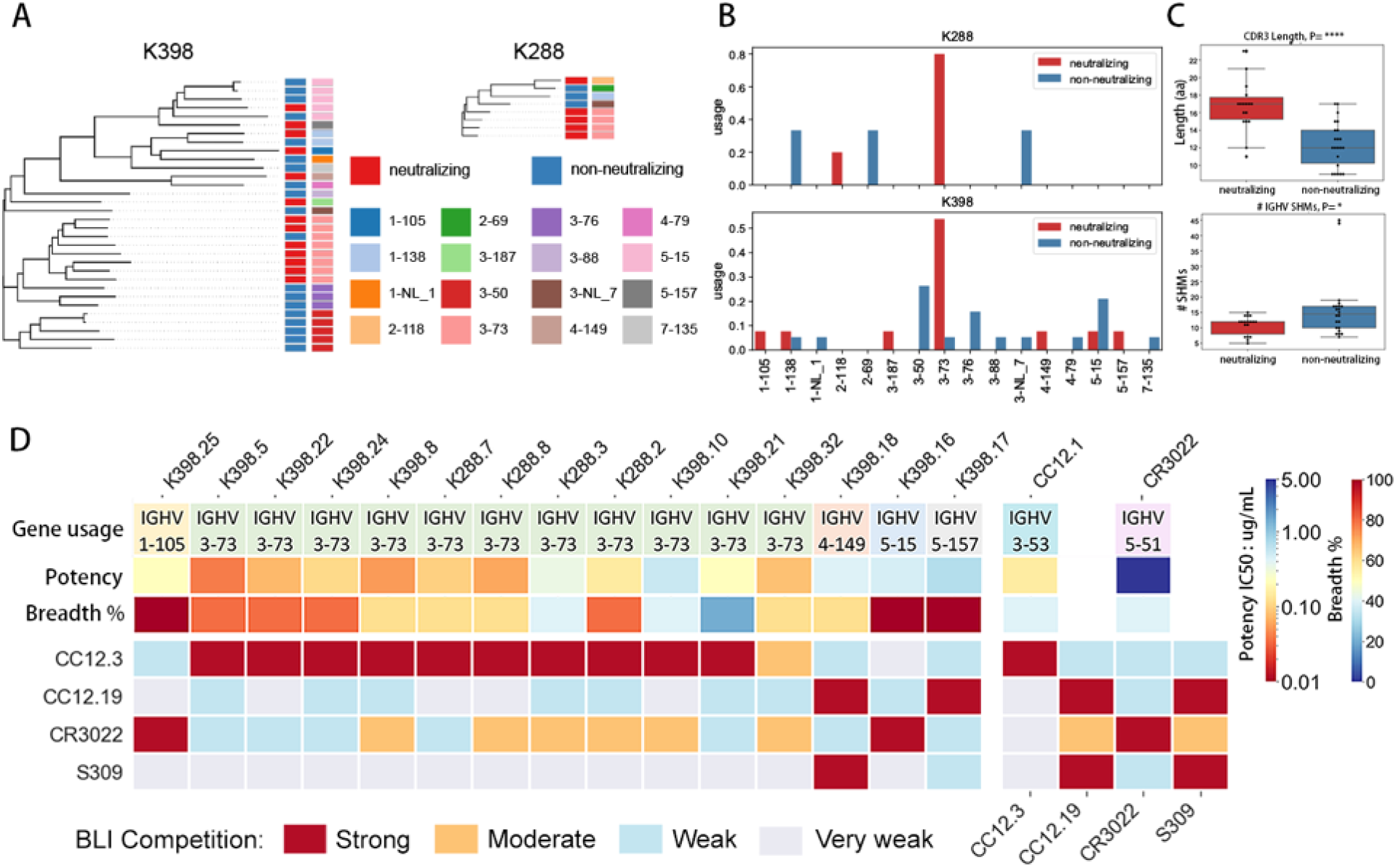
Immunogenetic and epitope properties of sarbecovirus cross-neutralizing macaque antibodies. **(A)** Phylogenetic trees derived from heavy chain (VH)-gene sequences of S-protein specific antibodies isolated from K398 and K288 macaques. Gene assignments used the rhesus macaque (*Macaca mulatta*) germline database described in (*62*). The IGHV gene usage for each mAb depicted in the tree is shown by different colors indicated in the lower panel of the color scheme. Neutralizing and non-neutralizing mAbs are shown in red and blue, respectively. **(B)** IGHV gene usage for neutralizing and non-neutralizing mAbs in K288 and K398 macaques. **(C)** CDRH3 length and number of SHMs for neutralizing and non-neutralizing mAbs. (*p <0.1; ****p < 0.0001). AA, amino acids. **(D)** Heatmap showing BLI competition-based epitope binning of macaque cross-neutralizing mAbs with human RBD-specific nAbs CC12.3, CC12.19, CR3022, and S309.The gene usage, geomean neutralization potency and breadth (calculated from neutralization with 5 viruses in figure 3B-C) for each nAb are indicated. The BLI competition experiment was performed with SARS-CoV-2 RBD, and the competition levels are indicated as red (strong), orange (moderate), light blue (weak) and grey (very weak competition).

We next mapped the epitopes recognized by the macaque RBD nAbs by BLI competition with CC12.3, CC12.19, CR3022 as well as S309 that recognizes the class 3 epitope site (*23, 24*). All 11 macaque IGHV3-73-encoded SARS-CoV-1/2 nAbs strongly competed with VH3-53 CC12.3 nAb, and many showed moderate to weak competition with CR3022 (Fig. 4D, fig. S12). Two nAbs, K398.17 and K398.18, strongly competed with CC12.19 nAb, with K398.18 antibody also competing with S309 (*47*). Two nAbs, K398.25 and K398.16 competed strongly with CR3022 that targets the class 4 site (Fig. 4D, fig. S12). The mAbs showing the greatest overall pan-sarbecovirus cross-neutralization breadth were those competing with CR3022 or CC12.19.

### Structural studies of macaque nAbs interacting with S-protein

We turned next to structural studies to gain a better understanding of the neutralizing cross-reactivity. Single-particle, negative-stain electron microscopy (nsEM) was used to image representative macaque nAb Fabs, K288.2, K398.22, K398.8 (VH3-73), K398.18 (VH4-149), K398.25 (VH1-105) and K398.16 (VH5-15) with SARS-CoV-1/2 S-proteins. NsEM confirmed binding of all 6 nAbs to spike RBD where the cross-nAbs encoded by various germline V-genes interacted with the RBD via distinct binding modes (Fig. 5A, fig. S13). The binding modes were largely similar for SARS-CoV-2 and SARS-CoV-1 S-proteins with some exceptions (Fig. 5, A and B, fig. S13). We compared reconstructions of two macaque IGVH3-73 and two human IGVH3-53 nAbs and noted differences in the angles of approach to the SARS-CoV-2 RBD; the macaque cross nAbs interacted with the RBD from a more lateral angle compared to the two human nAbs with a more perpendicular approach angle close to the spike 3-fold axis (Fig. 5C).

**Fig. 5.**
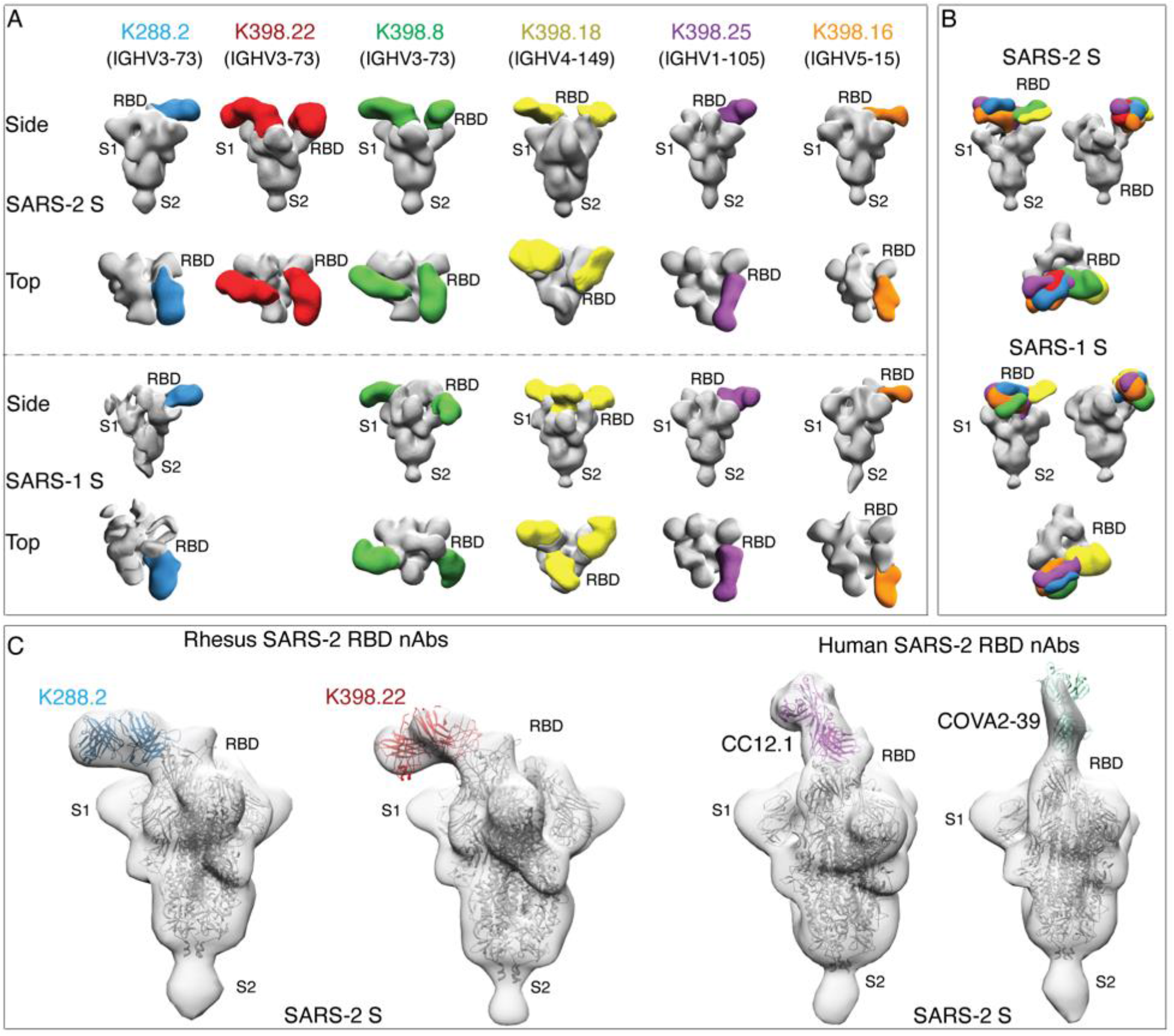
Electron microscopy structures of sarbecovirus cross-neutralizing macaque mAbs with SARS-CoV-2 and SARS-CoV-1 S-proteins. **(A)** Electron microscopy 3D reconstructions of rhesus macaque nAb Fabs with SARS-CoV-2 and SARS-CoV-1 S-proteins. Fabs of six RM nAbs, K288.2 (blue), K398.22 (red), K398.8 (green), (K398.18 (yellow), K398.25 (purple) and K398.16 (orange) were complexed with SARS-CoV-2 and SARS-CoV-1 S-proteins and 3D reconstructions were generated from 2D class averages. The heavy chain germline gene usage is indicated for each macaque nAb. The spikes S1 and S2 subunits and the RBD sites are labelled. **(B)** Side and top views showing 3D reconstructions of all 6 RBD-directed macaque cross-nAbs bound to SARS- CoV-2 (upper panel) and SARS-CoV-1 (lower panel) S-proteins. **(C)** S-protein binding angles of approach comparisons of macaque IGHV3-73-encoded RBD cross nAbs, K288.2 and K398.22, with human IGHV3-53-encoded RBD nAbs, CC12.1 and COVA2-39, isolated from COVID-19 donors. The crystal structures of the respective nAbs were docked into EM density maps. The RBD, S1 and S2 subunits of SARS-CoV-2 S protein are labelled. The macaque nAbs approach spike RBD from the side and the two human nAbs approach at a more perpendicular angle close to the spike 3-fold axis.

To investigate the binding of the potent macaque VH3-73 nAbs in greater detail, we determined crystal structures of K288.2 and K398.22 antibody Fabs with SARS-CoV-2 RBD at 2.3 Å and 1.95 Å resolution, respectively (Fig. 6, A to H, tables S1 and S2). Both nAbs bind a novel epitope where the heavy-chain dominates the interaction as expected from enrichment of IGVH3-73 and its pairing with different light chains (Fig 6, A to H, figs. S10, S14, S15). However, the IGHV3 -73- encoded macaque nAbs and IGHV3-53 human nAbs interact differently with the RBD; both interact with the ACE2 binding site but at opposite ends (Fig 6, B and F, fig. S14, B and C). For the human nAbs, the germline-encoded CDRH1 and CDRH2 both contribute substantially to RBD recognition (*18, 22*), the macaque nAbs predominantly rely on CDRH2 and less on CDRH1 (Fig 6, D, H and I, fig. S14, C to E). Further, these CDRs bind to different regions on the RBD (fig. S14C). Most of the CDRH2-mediated epitope recognition by the macaque bnAbs derives from germline residues that interact with a conserved sarbecovirus RBD region (Fig 6, I and K, fig. S1). The human and macaque germlines also differ at critical CDRH1 residues. The human IGHV3 - 53 germline encodes a CDRH1 ^32^NY^33^ motif critical for RBD recognition (fig. S14, C and D) (*22*), whereas macaque IGHV3-73 antibodies bind to a different RBD region where only V_H_ E33 in CDRH1 of K288.2 and K398.22 interact (Fig. 6D, H, fig. S14, C and E). Thus, immunodominant IGHV3-73-encoded macaque nAbs target more highly conserved RBD residues across hACE2- utilizing sarbecoviruses thereby providing a structural basis for their broad cross-reactivity (Fig. 6K, fig. S16).

**Fig. 6.**
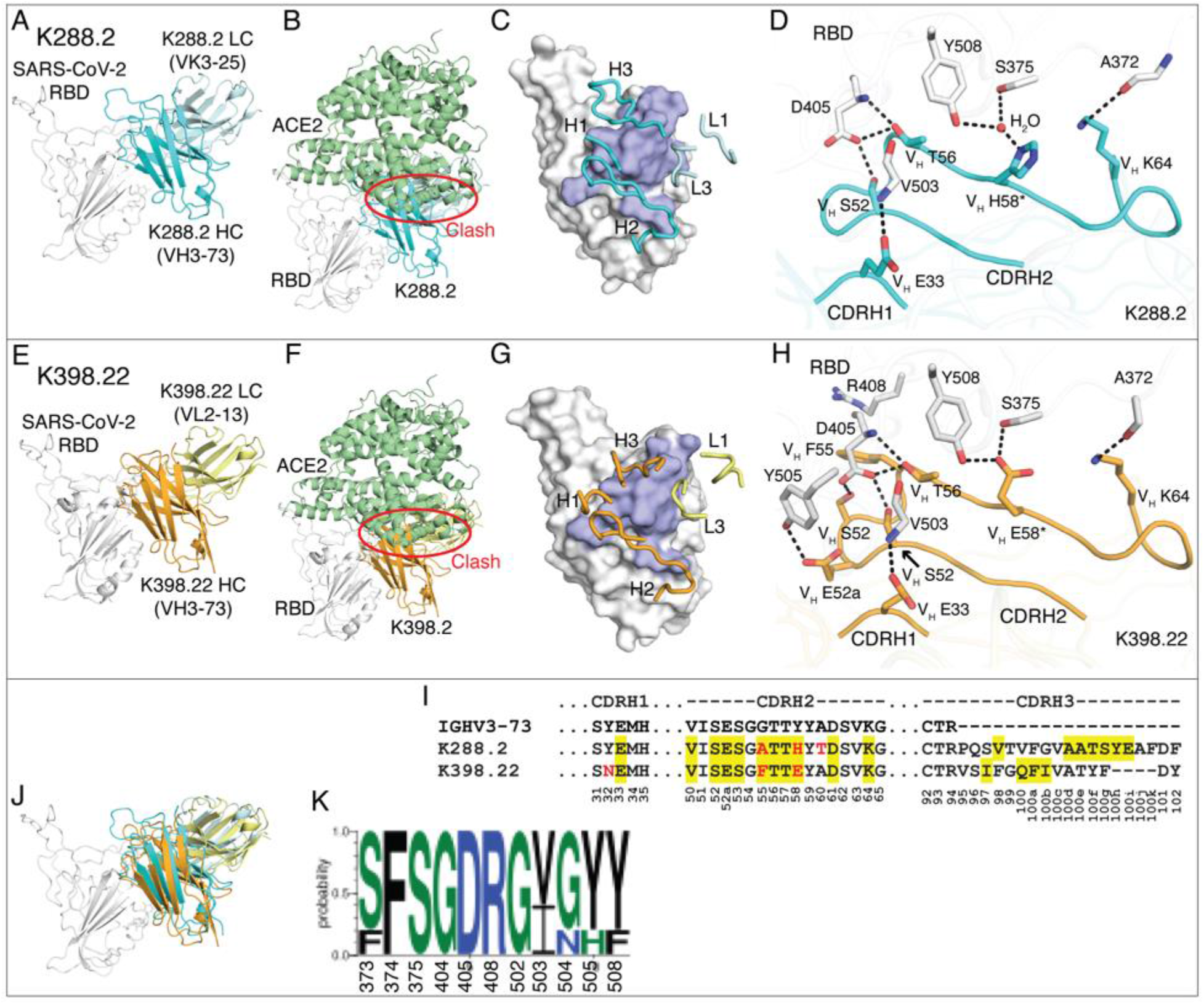
Crystal structures of two rhesus macaque IGHV3-73-encoded neutralizing antibodies bound to SARS-CoV-2 RBD. The SARS-CoV-2 RBD is shown in white throughout the figure, while human ACE2 (hACE2) is in pale green, heavy and light chains of K288.2 are in cyan and light cyan, and those of K398.22 in orange and yellow. For clarity, only variable domains of the antibodies are shown. Structures of SARS-CoV-2 RBD in complex with K288.2 antibody **(A-D)** and in complex with K398.22 antibody **(E-H)**. Germline genes encoding K288.2 and K398.22 macaque antibodies are labeled in **(A, E)**. Structures of SARS-CoV-2 RBD in complex with K288.2 or K398.22 superimposed onto an RBD/hACE2 complex structure (PDB 6M0J, (*74*)) show that the K288.2/K398.22 antibody would clash with the hACE2 receptor (indicated with a red circle). **(B, F)** The epitope of K288.2 or K398.22 is shown in purple. CDR loops that interact with the RBD are labeled **(C, G)**. **(D, H)** Detailed molecular interactions of K288.2 **(D)** or K398.22 **(H)** CDRs H1 and H2 with SARS-CoV-2 RBD. Somatically mutated residues are indicated with *. The CDRH2 of IGHV3-73 antibodies interacts extensively with the RBD. **(I)** IGHV3-73 germline sequence; black. Only CDR sequences are shown for clarity. Lengths of CDR H3 can vary. Yellow shades: paratope region in antibody heavy chain (BSA > 0 Å^2^). Red letters: somatic mutations. Bottom numbers: Kabat numbering. Y58 is mutated to E/H. Our structures show germline Y58 would clash, whereas E/H is able to interact with the RBD. **(J)** The Fab/RBD complex structures of the two antibodies are superimposed on the RBD. **(K)** Sequence conservation of the epitope residues in SARS-related strains recognized by IGHV3-73 encoded paratopes (cutoff = 4 Å). SARS-CoV-2, SARS-CoV-1, WIV-1, RaTG13, and pang17 are used for the conservation analysis.

## Discussion

Natural infection and current vaccines for SARS-CoV-2 in humans typically induce very limited cross-neutralizing responses. The emerging SARS-CoV-2 VOCs and the possibility of further SARS pandemics have increasingly focused attention on development of vaccines and immunization strategies that could induce potent cross-neutralizing antibody responses. Preclinical approaches to vaccine development rely heavily on immunization of animal models with candidate immunogens. Indeed, for the current COVID-19 vaccines, strong nAb responses were first observed to the SARS-CoV-2 S-protein presented on several platforms in a number of animal models (*38-40, 44, 48-55*). The nAb responses were protective in rhesus macaque models of SARS-CoV-2 infection (*5, 44, 49, 56, 57*) and clinical studies subsequently identified nAbs as a strong correlate of protection in humans (*58–60*). Most macaque studies focused on the potency of nAb responses to the original Wuhan or Washington strains of SARS-CoV-2 but, more recently, greater interest has centered on the breadth of responses to divergent viruses. One recent study (*49*) investigated use of different adjuvants with stabilized recombinant SARS-CoV-2 S proteins presented on nanoparticles in macaques and noted nAb responses of varying potency and neutralization breadth. One adjuvant produced only a 4.5-fold reduction in neutralizing titer against the B.1.351 variant, whereas another produced a 16-fold reduction. Another macaque study (*44*), which used RBD on a nanoparticle with a strong adjuvant as well as mRNA, induced very high nAb titers to SARS-CoV-2 with some reduction against VOCs. In addition, sera neutralized SARS-CoV-1 and related sarbecoviruses, albeit with reduced potency.

Here, we immunized macaques with S-protein using a strong saponin-based adjuvant and generated potent and broad nAb responses. Unlike other studies, nAb responses were equally potent against SARS-CoV-2 and SARS-CoV-1. Whereas the immunodominant human nAb RBS- A/Class 1 response to S-protein largely utilizes the VH3-53 germline gene segment, the immunodominant macaque response leads to a strong VH3-73 response that involves a different epitope and different approach angle to interact with the RBD. The different binding mode allows for SARS-CoV-1/SARS-CoV-2 cross-neutralization and retention of neutralization of VOCs.

A number of lessons can be learned from this study. First, one cannot necessarily assume that because immunization of macaques leads to SARS-CoV-2 cross-neutralizing responses that the same immunogen will give similar results in humans. Differences between human and rhesus macaque antibody repertoires can potentially have important consequences for vaccine model studies and this observation goes beyond SARS-CoV-2. For example, for HIV, VRC01-class broadly neutralizing antibodies are encoded by IGHV1-2 and no similar germlines are present in the macaque repertoire (*61–63*). An immunogen (eOD-GT8) specifically designed to activate human VH1-2 germline-encoded naive B cell precursors would then not be expected to produce a similar response in macaques. Second, the isolation of potent macaque cross-neutralizing antibodies furnished evidence of an epitope in SARS-CoV-2 that could be targeted by immunization of humans if the immunodominance of VH3-53 and other preferred germlines can be reduced by rational immunogen engineering (*64–67*). Although this particular epitope has not yet been targeted by human SARS-CoV-2 antibodies with known structures, the human SARS-CoV-2 nAb DH1047 (*68*) heavy chain binds in a similar location but in a completely different orientation so that the light chains are oppositely positioned (fig. S15). Third, we used a strong saponin-based adjuvant that may have contributed to elicitation of VH3-73 antibodies and adjuvants should be evaluated in combination with immunogens designed to induce pan-CoV nAbs, perhaps in small-scale experimental medicine studies, as emphasized elsewhere (*49*) Fourth, given the important response differences between human and the closely phylogenetically related rhesus macaque, continued exploration of human antibody-repertoire based model systems such as humanized mice (*69–71*) and specific antibody knock-in mice (*67, 72, 73*) should be a priority for immunogen evaluation in rational vaccine design approaches. These findings suggest caution in the use of macaque models in SARS vaccine evaluation but also provide valuable information that may ultimately assist in pan-CoV vaccine design.

## ACKNOWLEDGEMENTS

We thank all the human cohort participants for donating samples.

## Author contributions

W.H., M.Y., S.C., S.C., D.J.I., A.B.W., I.A.W., D.R.B. and R.A. conceived and designed the study. W.H., and S.C., G.S., D.H., M.F., and W.R. conducted animal immunization studies. M.S., and M.M. prepared the SMNP adjuvant. N.B., J.R., M.P., E.G., S.A.R., D.M.S., and T.F.R. recruited donors and collected and processed plasma samples. W.H., S.C., R.M., G.S., F.A., P.Y., and P.Z. performed BLI, ELISA, virus preparation, neutralization and isolation and characterization of monoclonal antibodies. F.Z. prepared the human antibodies. L.P., and D.H. prepared virus mutant plasmids. R.M., B.B., and Y.S. performed immunogenetic analysis of the antibodies. W.L., and J.T. conducted negative stain electron microscopy studies. M.Y. and X.Z. determined crystal structure of the antibody-antigen complex. W.H., M.Y., S.C., R.M., G.S., N.B., W.L., P.Y., J.T., P.Z., X.Z., L.P., D.H., F.A., D.N., B.B., Y.S., T.F.R., S.C., D.J.I., A.B.W., I.A.W., D.R.B., and R.A. designed the experiments and/or analyzed the data. W.H., M.Y., R.M., I.A.W., D.R.B. and R.A. wrote the paper, and all authors reviewed and edited the paper.

## Data and materials availability

The data supporting the findings of this study are available within the paper and its supplementary information files or from the corresponding author upon reasonable request. Antibody sequences have been deposited in GenBank under accession numbers XXX-XXX. The X-ray coordinates and structure factors will be deposited to the RCSB Protein Data Bank under accession code: XXXX. The EM maps will be deposited in the Electron Microscopy Data Bank (EMDB) under accession codes: YYYY

## Supplementary Materials

### MATERIALS AND METHODS

#### Human samples

##### Convalescent COVID-19 and SARS-CoV-2 spike vaccinated human sera

Sera from convalescent COVID-19 donors (*75*) and from spike-mRNA vaccinated humans were provided through the “Collection of Biospecimens from Persons Under Investigation for 2019- Novel Coronavirus Infection to Understand Viral Shedding and Immune Response Study” UCSD IRB# 200236. Protocol was approved by the UCSD Human Research Protection Program. Convalescent serum samples were collected based on COVID-19 diagnosis regardless of gender, race, ethnicity, disease severity, or other medical conditions. All human donors were assessed for medical decision-making capacity using a standardized, approved assessment, and voluntarily gave informed consent prior to being enrolled in the study.

#### Ethics Statement

The mice and rhesus macaque animal studies were approved and carried out in accordance with protocols provided to the Institutional Animal Care and Use Committee (IACUC) respectively at The Scripps Research Institute (TSRI; La Jolla, CA) under approval number 19-0020 and at Alphagenesis Institutional Animal Care and Use Committee (IACUC) under approval number AUP 19-10. The animals at both facilities were kept, immunized, and bled in compliance with the Animal Welfare Act and other federal statutes and regulations relating to animals and in adherence to the Guide for the Care and Use of Laboratory Animals (National Research Council, 1996).

#### Immunization and Sampling

8 outbred, healthy Indian-origin rhesus macaques (*Macaca mulatta*) were housed at Alphagenesis, Yemassee, SC. 2 groups of 4 rhesus macaques, evenly distributed by gender between the ages 3-5 years, were immunized with SARS-CoV-2 S protein immunogen as bolus prime (group-1) or escalating dose prime (group-2) and bolus immunization boost for both groups. Animals were primed (wk0) and boosted (wk10) by subcutaneously (SubQ) injecting 100µg of SARS-CoV-2 S-protein immunogen along with 375µg of a saponin adjuvant (SMNP) (Silva M. et. al., submitted) per animal per immunization. All immunizations were administered SubQ divided between the left and right mid-thighs. For escalating dose priming, animals were given seven injections of SARS-CoV-2 S protein and the SMNP adjuvant in each thigh over 12 days (on days 0, 2, 4, 6, 8, 10, 12). The total doses of SARS-CoV-2 S-protein immunogen at each injection during the escalating dose prime immunization were: 0.2, 0.43, 1.16, 3.15, 8.56, 23.3, 63.2ug distributed evenly between two immunization sites. EDTA Blood was collected at various time points into CPT tubes for PBMC and plasma isolation. Serum was isolated using serum collection tubes and frozen. One group of 5 mice (C57BL/6 mice (Jackson Laboratory): 3 females and 2 males), aged ∼8 weeks, were immunized with 20µg SARS-CoV-2 S protein immunogen along with 5µg of SMNP adjuvant per animal per immunization. Animals were immunized twice subcutaneously by infection immunogen-adjuvant vaccine formulation at prime (wk0) and boost (wk3) and the serum samples were collected at various time points for the analysis of antibody responses.

#### Plasmid construction

To generate the expression plasmids for soluble S ectodomain proteins from SARS-CoV-1 (residues 1-1190; GenBank: AAP13567) and SARS-CoV-2 (residues 1-1208; GenBank: MN908947), spike genes were synthesized by GeneArt (Life Technologies). The ectodomains of SARS-CoV-1 and SARS-CoV-2 were constructed by PCR amplification and Gibson assembly (NEB, E2621L) cloning into the vector phCMV3 (Genlantis, USA). To stabilize soluble S proteins in the trimeric prefusion state, we made the following changes: double proline substitutions (2P) in the S2 subunit, replacement of the furin cleavage site by “GSAS” in SARS-CoV-2 (residues 682–685), and SARS-CoV-1 (residues 664–667), and incorporation of a C-terminal T4 fibritin trimerization motif (*76, 77*). To aid purification and biotinylation, the HRV-3C protease cleavage site, 6x HisTag, and AviTag spaced by GS-linkers were added to the C-terminus. To generate gene fragments encoding SARS-CoV-1 RBD (residue 307-513), SARS-CoV-2 NTD (residue 1- 290), RBD (residue 320-527), RBD-SD1 (residue 320-591), and RBD-SD1-2 (residue 320-681) subdomains, PCR-amplifications were carried out from the SARS-CoV-1 and SARS-CoV-2 plasmids. The original secretion signal or the Tissue Plasminogen Activator (TPA) leader sequence were cloned in frame with the gene fragments.

#### Transient transfection

To express antibodies, the corresponding HC and LC plasmids were transiently transfected into the Expi293 cell (Life Technologies) at 3 x 10^6^ cells/mL with FectoPRO PolyPlus transfection reagent (Polyplus Cat # 116-040). To boost the protein yield, we fed cells with 300 mM sodium valproic acid solution and glucose solution 1 day post transfection. To purify monoclonal antibodies, the supernatants were harvested 4 days post transfection. To express the soluble S ectodomain proteins from SARS-CoV-1, SARS-CoV-2 and their truncations, plasmids were transfected into HEK293F cells (Life Technologies) at 1 million cells/mL. For 1L HEK293F transfection, we combined 350 μg plasmids with 16 mL transfectagro™ (Corning) and filtered with 0.22 μm Steriflip™ Sterile Disposable Vacuum Filter Units (MilliporeSigma™). The 1.8 mL 40K PEI (1 mg/mL) was mixed into 16 mL transfectagro™ in another Conical tube. The filtered plasmid solution was gently combined with the mixed PEI solution by inverting the tube several times. After resting at room temperature for 30 mins, the mixture was poured into 1L HEK293F cells. The supernatant was centrifuged and filtered with 0.22 μm membrane to a glass bottle 5 days post transfection.

#### Protein purification

To purify the mAbs, Expi293 supernatant was loaded on to Protein A Sepharose (GE Healthcare) for 2h at room temperature or 4°C overnight. After incubation, the Protein A Sepharose was loaded into Econo-Pac columns (BioRad #7321010). 1 column volume of PBS was used to wash away nonspecific protein. To elute the antibody, 0.2 M citric acid (pH 2.67) was briefly mixed with Protein A Sepharose and then dripped into 2M Tris Base to neutralize the protein solution. The antibody solution was buffer exchanged into PBS using 30K Amicon tubes (Millipore, UFC903008). To purify the His-tagged proteins, HEK293F supernatant were loaded on to HisPur Ni-NTA Resin (Thermo Fisher). Three bed volumes of wash buffer (25 mM Imidazole, pH 7.4) were used to wash away the nonspecific binding proteins before the elution step. To elute the purified proteins from the column, 25 mL of elution buffer (250 mM Imidazole, pH 7.4) was loaded to the column at slow gravity speed (∼4 sec/drop). The purified proteins were buffer exchanged into PBS using Amicon tubes. For further purification, the proteins were subjected to size exclusion chromatography using Superdex 200 (GE Healthcare). Endotoxin from the SARS-CoV- 2 spike protein immunogen was removed using Pierce™ High-Capacity Endotoxin Removal Resin (#88270) following manufacturer’s directions. Briefly, 50 µL of beads were washed twice and resuspended in 1 mL of filtered PBS. 500 µL of protein (2 mg/mL) was added to the resin and incubated at 4°C overnight on a rocking incubator. Protein and beads were passed through a 0.22 µm spin filter. Endosafe nexgen-MCS (Charles River) and Endosafe LAL cartridges (Charles River #PTS201F) were used to measure endotoxin levels. An endotoxin level below 10 EU/mL was used for the immunizations.

#### Cell lines

To generate HeLa-hACE2 cells, the human ACE2 lentivirus was transduced into HeLa cells. To produce the ACE2 lentivirus, the pBOB-hACE2 plasmid was co-transfected into HEK293T cells with lentiviral packaging plasmids pMDL, pREV, and pVSV-G (Addgene #12251, #12253, #8454) by Lipofectamine 2000 (Thermo Fisher Scientific, 11668019). The supernatants that contain lentivirus were collected 48 h post transfection. The stable cell line post transduction was selected and scaled up for use in the neutralization assay. HEK293F cells is a human embryonic kidney suspension cell line (Life Technologies). We used 293FreeStyle expression medium (Life Technologies) to culture the cell line in the shaker at 150 rpm, 37°C with 8% CO_2_. Expi293F cells are derived from the HEK293F cell line (Life Technologies). We used Expi293 Expression Medium (Life Technologies) in the same condition as the HEK293F cell line. The adherent cells were cultured in flasks or dishes in the incubator at 37°C with 8% CO_2_ with DMEM (vendor, Cat #) supplemented with 10% FBS (Vendor, Cat #) and 1% penicillin-streptomycin.

#### ELISA

96-well half-area plates (Corning cat. #3690, Thermo Fisher Scientific) were coated overnight at 4°C with 2 μg/mL of mouse anti-His-tag antibody (Invitrogen cat. #MA1-21315-1MG, Thermo Fisher Scientific) in PBS. Plates were washed 3 times with PBS plus 0.05% Tween20 (PBST) and blocked with 3% (wt/vol) bovine serum albumin (BSA) in PBS for 1 h. After removal of the blocking buffer, the plates were incubated with His-tagged spike proteins at a concentration of 5 μg/mL in 1% BSA plus PBST for 1.5 h at room temperature. After a washing step, NHP serum samples were added in 3-fold serial dilutions in 1% BSA/PBST starting from 1:30 dilution, and incubated for 1.5 h. For positive controls, CR3022 was used for SARS-CoV-1 spike, CC12.1 for SARS-CoV- 2 spike, and Lotus 6C serum for MERS-CoV spike. DEN3 was used as a negative control for all 3 spike proteins. Control antibodies were added in 3-fold serial dilutions in 1% BSA/PBST starting at 10 μg/mL. After washes, a secondary antibody conjugated with alkaline phosphatase (AffiniPure goat anti-human IgG Fc fragment specific, Jackson ImmunoResearch Laboratories cat. #109-055-008) diluted 1:1000 in 1% BSA/PBST, was added to each well. After 1 h of incubation, the plates were washed and developed using alkaline phosphatase substrate pNPP tablets (Sigma cat. #S0942-200TAB) dissolved in a stain buffer. The absorbance was measured after 10 and 20 minutes and was recorded at an optical density of 405 nm (OD405) using a VersaMax microplate reader (Molecular Devices), where data were collected using SoftMax version 5.4 (Molecular Devices). The wells without the addition of serum served as a background control.

#### Pseudovirus production

To generate the pseudoviruses, the MLV-gag/pol and MLV-CMV-Luciferase plasmids were co-transfected with full-length or variant SARS-CoV-1 or SARS-CoV-2 plasmid into HEK293T cells by using Lipofectamine 2000 (Thermo Fisher Scientific, 11668019). The supernatants containing pseudovirus were collected 48 h post transfection and filtered by a 0.22 μm membrane. The pseudovirus was stored at -80°C prior to use.

#### Pseudovirus entry and serum neutralization assays

To test the inhibition of pseudovirus infection by serum or mAbs, we used the stable cell line HeLa-hACE2 generated by lentivirus transduction with consistent ACE2 expression level to carry out the assay. To calculate the neutralization efficiency, the sera or mAbs were 3-fold serially diluted. 25 μL of each dilution was incubated with 25 μL of pseudovirus at 37 °C for 1 h in 96-half area well plates (Corning, 3688). During the incubation time, HeLa-hACE2 cells were suspended at a concentration of 2 x 10^5^/mL with the culture medium adding DEAE-dextran (20 μg/mL, Sigma, # 93556-1G). After 1 h incubation, 50 μL of cell solution was distributed into each well (10,000 cells/well). To evaluate the neutralization efficiency, the supernatants were removed from the well 48 h post infection. The HeLa-hACE2 cells were lysed by luciferase lysis buffer (25 mM Gly-Gly pH 7.8, 15 mM MgSO_4_, 4 mM EGTA, 1% Triton X-100) at room temperature for 10-20 mins. Luciferase activity of the cells in each well were inspected after adding Bright-Glo (Promega, PR- E2620) by a luminometer. All of the sera and mAbs were tested in duplicate for each experiment. All experiments were repeated independently at least twice. Percentage of neutralization was calculated according to the equation:

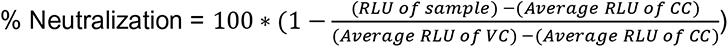

The 50% pseudovirus neutralizing (IC_50_) or binding (ID_50_) antibody titer was calculated by non-linear fitting the plots of luciferase signals against antibody concentrations or sera dilution ratio in Graph Pad Prism.

For neutralization of anti-RBD antibody depleted macaque sera, the immune sera were depleted by two rounds of sequential incubation with SARS-CoV-2 RBD-conjugated with magnetic beads. Beads were conjugated to the RBD by mixing every 5 mg of Streptavidin T1 beads (Invitrogen™ Dynabeads™ MyOne™ Streptavidin T1, Catalog number: 65601) with 75 µg of randomly biotinylated SARS-CoV-2-RBD protein. Prior to conjugation, 5 mg of Streptavidin T1 beads were washed with 1 mL of 0.1% BSA in 1x PBS 4 times and once with 1 mL of 1x PBS. A DynaMag™- 2 Magnet was used to capture the beads during washes. For conjugation, 500 µL (500 µg/mL) of randomly biotinylated SARS-CoV-2-RBD solution was mixed with 5 mg of cleaned T1 beads. The mixture was placed on a rotator at room temperature for 1 hour. To wash away unconjugated SARS-CoV-2-RBD protein, the sample was washed with 1x PBS 3 times. To deplete the RBD-binding antibodies from the immune sera, 60 µl of each serum sample was diluted 5-fold and mixed with 5 mg of the RBD-conjugated magnetic beads and incubated overnight at 4℃. Another round of depletion was done using 5 mg of the RBD-conjugated magnetic beads at 4℃ for 3 hours to further eliminate any remaining RBD-binding antibodies. The depletion of RBD directed antibodies in the immune sera was confirmed by BLI binding to SARS-CoV-2 RBD and the S-protein.

#### Bio-layer interferometry (BLI)

Binding of NHP mAbs to SARS-CoV-1, SARS-CoV-2 spike proteins and truncated proteins was analyzed on an Octet RED384 system using a Protein A biosensor (18-5010, Sartorius) to capture the mAbs. 10 μg/mL of each NHP mAbs were loaded on the hydrated biosensor for 60s. Biosensors were subsequently moved into blank buffer (1x PBS + 0.1% Tween 20) for 60s to remove unbound protein and provide a baseline. The biosensors were moved to immerse in the protein solutions with 200 nM spike/truncated proteins for 120s to acquire an association signal. To monitor disassociation, the biosensors were moved into blank buffer for 240s. All proteins and mAbs were diluted with 1x PBS + 0.1% Tween 20.

#### In-tandem epitope binning by BLI

In-tandem epitope binning was carried out using the Octet RED384 in order to distinguish binding epitopes of mAbs from each other or from human mAbs or from hACE2-Fc. 100 nM of His-tagged SARS-CoV2-RBD protein antigens were captured using Anti-HIS (HIS2) biosensors (18-5114, Sartorius). The biosensor was loaded with antigen for 5 min and then moved into the mAbs at saturating concentration of 100 µg/mL for 10 min. The biosensors were then moved into competing mAbs at a concentration of 25 µg/mL for 5 min to measure binding in the presence of saturating antibodies. All reactions were done in 1x PBS + 0.1% Tween 20.

#### Competition BLI

To inspect the binding epitope of the NHP sera compared with known human SARS-CoV-2 mAbs or hACE2-FC, we did in-tandem epitope binning experiments using the Octet RED384 system. 200 nM of randomly biotinylated SARS-CoV-2 S or RBD protein antigens were captured using SA biosensors (18-5019, Sartorius). The biosensor was loaded with antigen for 5 min and then moved into the saturating mAbs at a concentration of 100 µg/mL for 10 min. The biosensors were then moved into 1:50 diluted NHP sera for 5 min to measure binding in the presence of saturating antibodies. As control, biosensors loaded with antigen were directly moved into 1:50 diluted NHP sera.

The percent (%) inhibition in binding is calculated with the formula: [Percent (%) binding inhibition = 1- (serum binding response in presence of the competitor antibody / binding response of the corresponding control serum antibodies without the competitor antibody)]

#### Random biotinylation of proteins

The spike proteins and truncated proteins were randomly biotinylated using an EZ-Link NHS-PEG Solid-Phase Biotinylation Kit (Thermo Scientific #21440). The reagent in each tube were dissolved with 10 μL DMSO into stock solution. For the working solution, 1 μL stock solution was diluted by 170 μL water freshly before use. The spike protein or truncated proteins were concentrated to 7- 9 mg/mL using Amicon tubes in PBS before biotinylation. 3 μL of working solution was added into each 30 μL protein aliquot and incubated on ice for 3 h. The reaction ended by buffer-exchanging the protein into PBS. The biotinylated proteins were evaluated by BioLayer Interferometry.

#### BirA biotinylation of proteins for B cell sorting

To generate biotinylated spike probes for B cell sorting, the coronavirus spike was constructed with an Avi-tag at the C-terminus. The Avi-tagged spike proteins were concentrated to 7-9 mg/mL in TBS before the biotinylation reaction using BirA Biotin-Protein Ligase Reaction Kit (Avidity) following manufacturer’s instructions. Briefly, for each reaction, 50 μL of protein solution, 7.5 μL of BioB Mix, 7.5 μL of Biotin200, and 5 μL of BirA ligase (3 mg/mL) were added in PCR tubes and mixed thoroughly. After incubating on ice for 3 h, the biotinylated protein was purified by size-exclusion chromatography. The biotinylated proteins were evaluated by BioLayer Interferometry using the SA biosensor.

#### Isolation of monoclonal antibodies (mAbs)

The sorting strategy of antigen-specific memory B cells was described in previous papers (*78–80*). The sorting process was performed in a 96-well format. To enrich for antigen-specific memory B cells, PBMCs from NHP 14 weeks post immunization were stained with fluorescently labeled antibodies as well as the fluorophore-conjugated spike proteins. Specifically, BirA biotinylated SARS-CoV-1 spike was coupled to streptavidin-AF647 (Thermo Fisher S32357). The BirA biotinylated SARS-CoV-2 spike was coupled to streptavidin-AF488 (Thermo Fisher S32354), streptavidin-BV421 (BD Biosciences 563259) separately. The spike proteins were freshly conjugated with the streptavidin-fluorophores at 2:1or 4:1 molecular ratio at room temperature for 30 min before use. The frozen NHP PBMCs were thawed in 10 mL recover medium (RPMI 1640 medium containing 50% FBS) immediately before staining. The cells were washed with FACS buffer (PBS, 2% FBS, 2 mM EDTA) and counted. Each 10 million cells were resuspended in 100 μL of FACS buffer and labeled with antibodies for rhesus macaque surface markers. The T cell markers CD3 (APC Cy7, BD Pharmingen #557757), CD4 (APC-Cy7, Biolegend, #317418), CD8 (APC-Cy7, BD Pharmingen #557760); the monocytes marker CD14 (APC-H7, BD Pharmingen #561384, clone M5E2); and the IgM (PE, Biolegend, #314508, clone MHM-88) were stained for negative selection. The B cell markers CD19 (PerCP-Cy5.5, Biolegend, #302230, clone HIB19), CD20 (PerCP-Cy5.5, Biolegend, #302326, clone 2H7) and IgG (BV786, BD Horizon, #564230, Clone G18-145) were stained to select IgG+ B cells. After 15 min staining on ice, SARS-CoV-1- S-AF647, SARS-CoV-2-S-AF488, and SARS-CoV-2-S-BV421 were added to the staining solution. After another 30 min incubation on ice, FVS510 Live/Dead stain (Thermo Fisher Scientific, #L34966) 1:1000 diluted with FACS buffer was added into the staining solution. 15 min later, the stained cells were washed with cold 10 mL FACS buffer and resuspended in 500 μL FACS buffer for each 10 million cells. After filtering the cells through a 70 μm mesh cap FACS tube (Fisher Scientific, #08-771-23), the cells were sorted using a BD FACSMelody (BRV 9 Color Plate 4way). To isolate the SARS-CoV-1-S and SARS-CoV-2-S cross-reactive B cells, the gating strategy was designed as follows: after gating the lymphocytes (SSC-A vs. FSC-A) and singlets (FSC-H vs. FSC-A), live cells were selected by negative gating of FVS510 Live/Dead staining. The CD3, CD4, CD8, CD14 negative and CD19, CD20 positive cells were gated as “CD19/CD20+, Dump-”. By selecting the IgG positive and IgM negative cells, the cells in the “IgG^+^, IgM^-^” gate were sequentially selected for CoV-2-S-BV421/CoV-2-S-AF488 double positive and CoV-1-S-AF647/CoV-2-S-AF488 double positive reactivity. The selected cells were sorted into 96-well plates individually and plates moved onto dry ice immediately after sorting. cDNA was generated from the sorted cells on the same day to avoid degradation of RNA. The Superscript IV Reverse Transcriptase (Thermo Fisher), dNTPs (Thermo Fisher), random hexamers (Gene Link), Ig gene-specific primers, DTT, and RNAseOUT (Thermo Fisher), were used in the lysis buffer containing Igepal (Sigma) to carry out reverse transcription PCR as described previously (23). To amplify IgG heavy and light chain variable regions, cDNA of each single cell was used as a template in two rounds of nested PCR reactions using Hot Start DNA Polymerases (Qiagen, Thermo Fisher) and rhesus macaque primers as described previously (*80*). The PCR products were purified with SPRI beads according to manufacturer’s instructions (Beckman Coulter). Purified DNA fragments of heavy and light chain variable regions were subsequently cloned into expression vectors encoding human IgG1, and Ig kappa/lambda constant domains, respectively, using Gibson assembly (NEB, E2621L) according to the manufacturer’s instructions. Paired heavy and light chain were sequenced and analyzed using the rhesus macaque (*Macaca mulatta*) germline database from (*62*). The paired plasmids were co-transfected into 293Expi cells for antibody expression.

#### Fab production

To generate the Fab of an IgG, a stop codon was inserted in the heavy chain constant region at “KSCDK*”. The truncated heavy chains were co-transfected with the corresponding light chains in 293Expi cells to produce the Fab. The supernatants were harvested 4 days post transfection. Fabs were purified with CaptureSelect™ CH1-XL MiniChrom Columns (#5943462005). Supernatants were loaded onto columns using an Econo Gradient Pump (Bio-Rad #7319001) Following a wash with 1x PBS, Fabs were eluted with 25 mL of 50 mM acetate (pH 4.0) and neutralized with 2 M Tris Base. The eluate was buffer exchanged with 1x PBS in 10K Amicon tubes (Millipore, UFC901008) and filtered with a 0.22 µm spin filter.

#### Negative stain electron microscopy

Spike protein was complexed with Fab at three times molar excess per trimer and incubated at room temperature for 30 mins. Complexes were diluted to 0.02mg/ml in 1x Tris-buffered saline and 3µl applied to a 400mesh Cu grid, blotted with filter paper, and stained with 2% uranyl formate.

Micrographs were collected on a ThermoFisher Tecnai Spirit microscope operating at 120kV with a FEI Eagle CCD (4k) camera at 52,000 magnification using Leginon automated image collection software (*81*). Particles were picked using DogPicker (*82*) and 3D classification was done using Relion 3.0 (*83*).

#### Crystal structure determination of Fab-RBD complexes

The coding sequence for receptor binding domain (RBD; residues 333-529) of the SARS-CoV-2 spike (S) protein was synthesized and cloned into a customized pFastBac vector (*84*), which was designed to fuse an N-terminal gp67 signal peptide and C-terminal His_6_-tag to the target protein. To express the RBD protein, a recombinant bacmid DNA was generated from the sequencing-confirmed pFastBac construct using the Bac-to-Bac system (Life Technologies). Baculovirus was generated by transfecting purified bacmid DNA into Sf9 cells using FuGENE HD (Promega), and subsequently used to infect a suspension cultures of High Five cells (Life Technologies) at a multiplicity of infection (MOI) of 5 to 10. Infected High Five cells were incubated at 28 °C with shaking at 110 rpm for 72 hours for protein expression. RBD protein that was secreted into the supernatant was then concentrated using a 10 kDa MW cutoff Centramate cassette (Pall Corporation). The RBD protein in the concentrate was purified by affinity chromatography using Ni-NTA resin (QIAGEN), followed by size exclusion chromatography on a HiLoad Superdex 200 pg column (GE Healthcare), and buffer exchanged into 20 mM Tris-HCl pH 7.4 and 150 mM NaCl using the same protocol as before (*85*). Crystallization trials were set up for SARS-CoV-2 RBD in complex with K288.2 and K398.22 Fabs. The Fab/RBD complexes were formed by mixing the two components in an equimolar ratio and incubating overnight at 4°C before setting-up crystal trials. The complexes (13 mg/ml) were screened for crystallization using the 384 conditions of the JCSG Core Suite (Qiagen) on our high-throughput robotic CrystalMation system (Rigaku) at Scripps Research by the vapor diffusion method in sitting drops containing 0.1 μl of protein and 0.1 μl of reservoir solution. For the K288.2/RBD complex, optimized crystals were then grown in 0.1 M sodium cacodylate pH 6.5, 40% (v/v) 2-methyl-2,4-pentanediol, and 5% (w/v) polyethylene glycol (PEG) 8000 at 20°C. Crystals were grown for 7 days and then flash cooled in liquid nitrogen. Diffraction data were collected at cryogenic temperature (100 K) at the Stanford Synchrotron Radiation Lightsource (SSRL) on the Scripps/Stanford beamline 12-1 with a wavelength of 0.9793 Å, and processed with HKL2000 (*86*). For the K298.22/RBD complex, optimized crystals were then grown in 0.2 M ammonium formate and 20% (w/v) PEG 3350 at 20°C. Crystals were grown for 7 days, pre-equilibrated with 10% (v/v) ethylene glycol, and then flash cooled in liquid nitrogen. Diffraction data were collected at cryogenic temperature (100 K) at beamline 23ID-D of the Advanced Photon Source (APS) at the Argonne National Laboratory with a wavelength of 0.9793 Å, and processed with XDS (*87*). Structures were solved by molecular replacement using PHASER (*88*) with PDB 7LOP for the RBD (*23*), whereas the models for the Fabs were generated by Repertoire Builder (https://sysimm.ifrec.osaka367 u.ac.jp/rep_builder/) (*89*). Iterative model building and refinement were carried out in COOT (*90, 91*), respectively.

#### Phylogenetic analysis

Heavy chain sequences of neutralizing and non-neutralizing antibodies collected from animals K398 and K288 were processed using DiversityAnalyzer tool (*92*). IGHV, IGHD, and IGHJ genes of rhesus monkey (*Macaca mulatta*) described in (*62*) were used as the database of germline immunoglobulin genes. For each heavy chain sequence, the number of SHMs was computed as the number of differences in the V segment with respect to the closest IGHV gene from the database. Phylogenetic trees derived from heavy chain sequences of antibodies collected from animals K398 and K288 were constructed using ClusterW2 tool (*93*) and visualized using Iroki tool (*94*).

#### Statistical Analysis

Statistical analysis was performed using Graph Pad Prism 8 for Mac, Graph Pad Software, San Diego, California, USA. Group of data were compared using the non-parametric unpaired Mann-Whitney-U test. Data were considered statistically significant at p < 0.05.

---

**Table S1.**
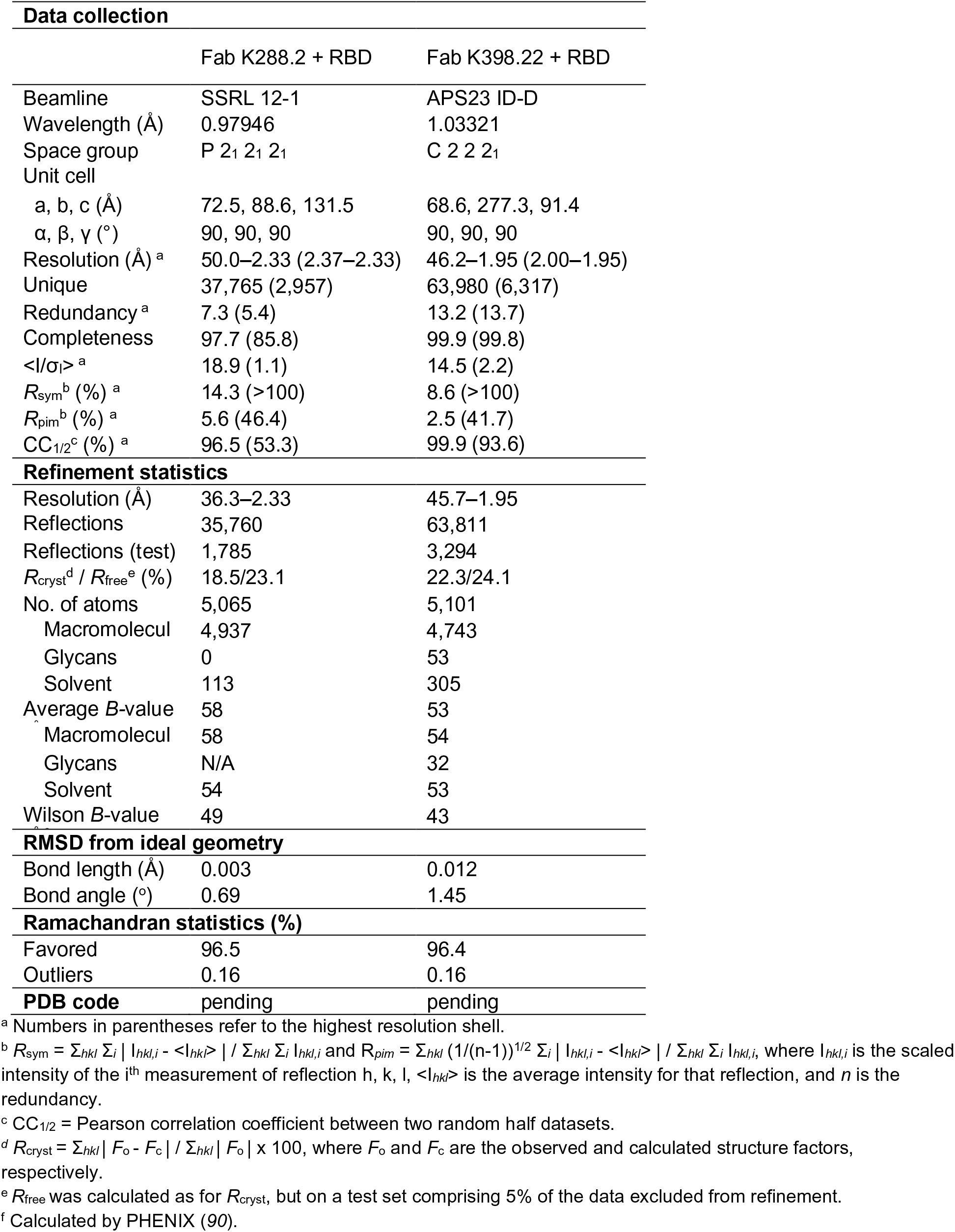
X-ray data collection and refinement statistics.

**Table S2.**
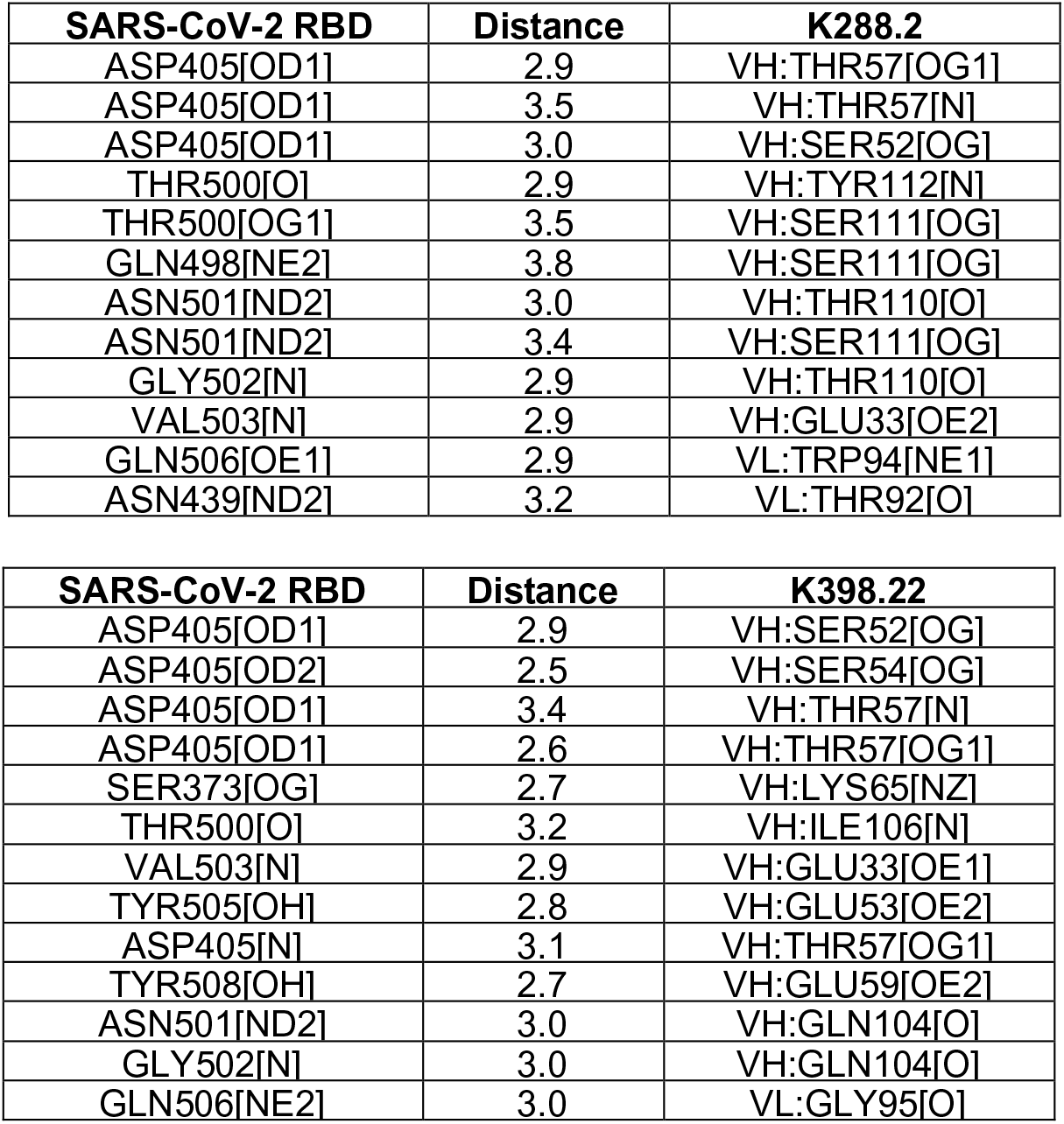
Hydrogen bonds identified at the antibody-RBD interface using the PISA program.

| **SARS-CoV-2 RBD** | **Distance** | **K288.2** |
| --- | --- | --- |
| ASP405[OD1] | 2.9 | VH:THR57[OG1] |
| ASP405[OD1] | 3.5 | VH:THR57[N] |
| ASP405[OD1] | 3.0 | VH:SER52[OG] |
| THR500[O] | 2.9 | VH:TYR112[N] |
| THR500[OG1] | 3.5 | VH:SER111[OG] |
| GLN498[NE2] | 3.8 | VH:SER111[OG] |
| ASN501[ND2] | 3.0 | VH:THR110[O] |
| ASN501[ND2] | 3.4 | VH:SER111[OG] |
| GLY502[N] | 2.9 | VH:THR110[O] |
| VAL503[N] | 2.9 | VH:GLU33[OE2] |
| GLN506[OE1] | 2.9 | VL:TRP94[NE1] |
| ASN439[ND2] | 3.2 | VL:THR92[O] |

| **SARS-CoV-2 RBD** | **Distance** | **K398.22** |
| --- | --- | --- |
| ASP405[OD1] | 2.9 | VH:SER52[OG] |
| ASP405[OD2] | 2.5 | VH:SER54[OG] |
| ASP405[OD1] | 3.4 | VH:THR57[N] |
| ASP405[OD1] | 2.6 | VH:THR57[OG1] |
| SER373[OG] | 2.7 | VH:LYS65[NZ] |
| THR500[O] | 3.2 | VH:ILE106[N] |
| VAL503[N] | 2.9 | VH:GLU33[OE1] |
| TYR505[OH] | 2.8 | VH:GLU53[OE2] |
| ASP405[N] | 3.1 | VH:THR57[OG1] |
| TYR508[OH] | 2.7 | VH:GLU59[OE2] |
| ASN501[ND2] | 3.0 | VH:GLN104[O] |
| GLY502[N] | 3.0 | VH:GLN104[O] |
| GLN506[NE2] | 3.0 | VL:GLY95[O] |

**Fig. S1.**
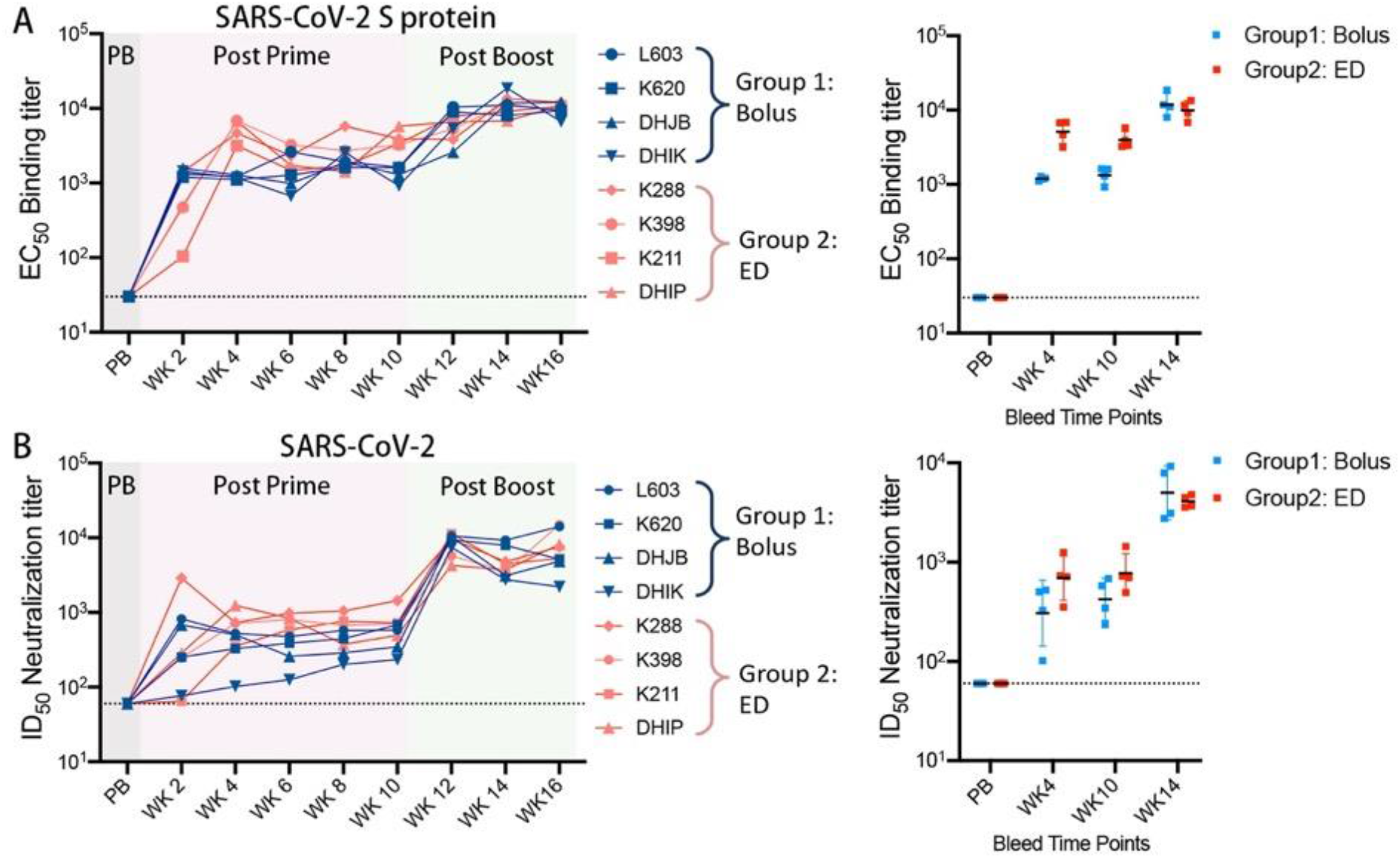
SARS-CoV-2 S protein prime-boost immunization in rhesus macaques induces strong antibody binding and neutralizing responses. **(A)** EC_50_ ELISA binding of the SARS-CoV-2 S protein prime (wk 2, 4, 6, 8, 10) and boosted (wk 12, 14 and 16) serum Ab responses to SARS-CoV-2 S protein in bolus (Group 1: blue) and ED (Group 2: red) immunization groups. PB = pre-bleed serum collected pre-immunization at wk0. Dot plots showing EC_50_ binding comparisons between bolus and ED groups at 4 time points (PB), post-prime (wk4 and wk10) and post-boost (wk14) immunizations time points. **(B)** SARS-CoV-2 virus-specific serum ID_50_ neutralizing antibody titers in SARS-CoV-2 S protein prime (wk 2, 4, 6, 8, 10) and boosted (wk 12, 14 and 16) animals in bolus (Group 1: blue) and ED (Group 2: red) immunization groups. PB = pre-bleed serum collected pre-immunization at wk0. Dot plots showing comparison of ID_50_ neutralization titers between bolus and ED groups at 4 time points (PB), post-prime (wk4 and wk10) and post-boost (wk14) immunization time points.

**Fig. S2.**
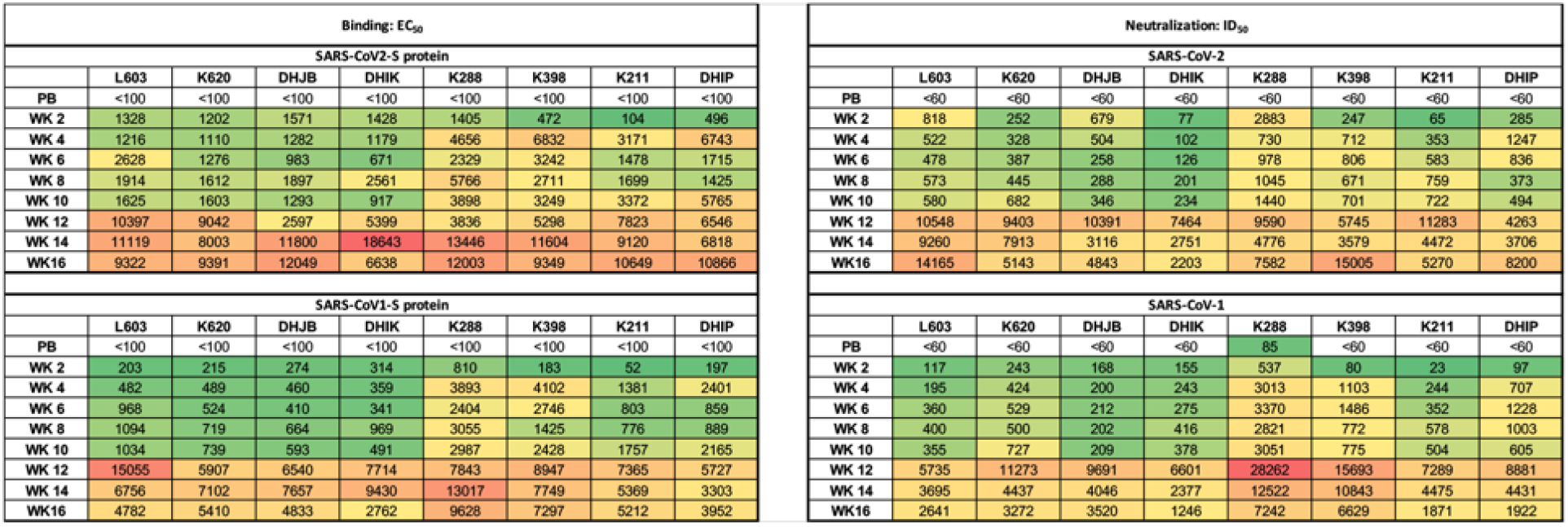
Binding and neutralization of rhesus macaque immune sera. SARS-CoV-2 S protein prime-boost immunization in rhesus macaques induces strong antibody binding and neutralizing responses against SARS-CoV-2 and cross-reactive binding and neutralizing responses against SARS-CoV-1. EC_50_ ELISA binding of the SARS-CoV-2 S protein prime (wk 2, 4, 6, 8, 10) and boosted (wk 12, 14 and 16) serum Ab responses to SARS-CoV-2 and SARS-CoV-1 S-proteins are shown on the left. SARS-CoV-2 and SARS-CoV-1 specific serum ID_50_ neutralizing antibody titers in SARS-CoV-2 S protein prime (wk 2, 4, 6, 8, 10) and boosted (wk 12, 14 and 16) animals are shown on the right.

**Fig. S3.**
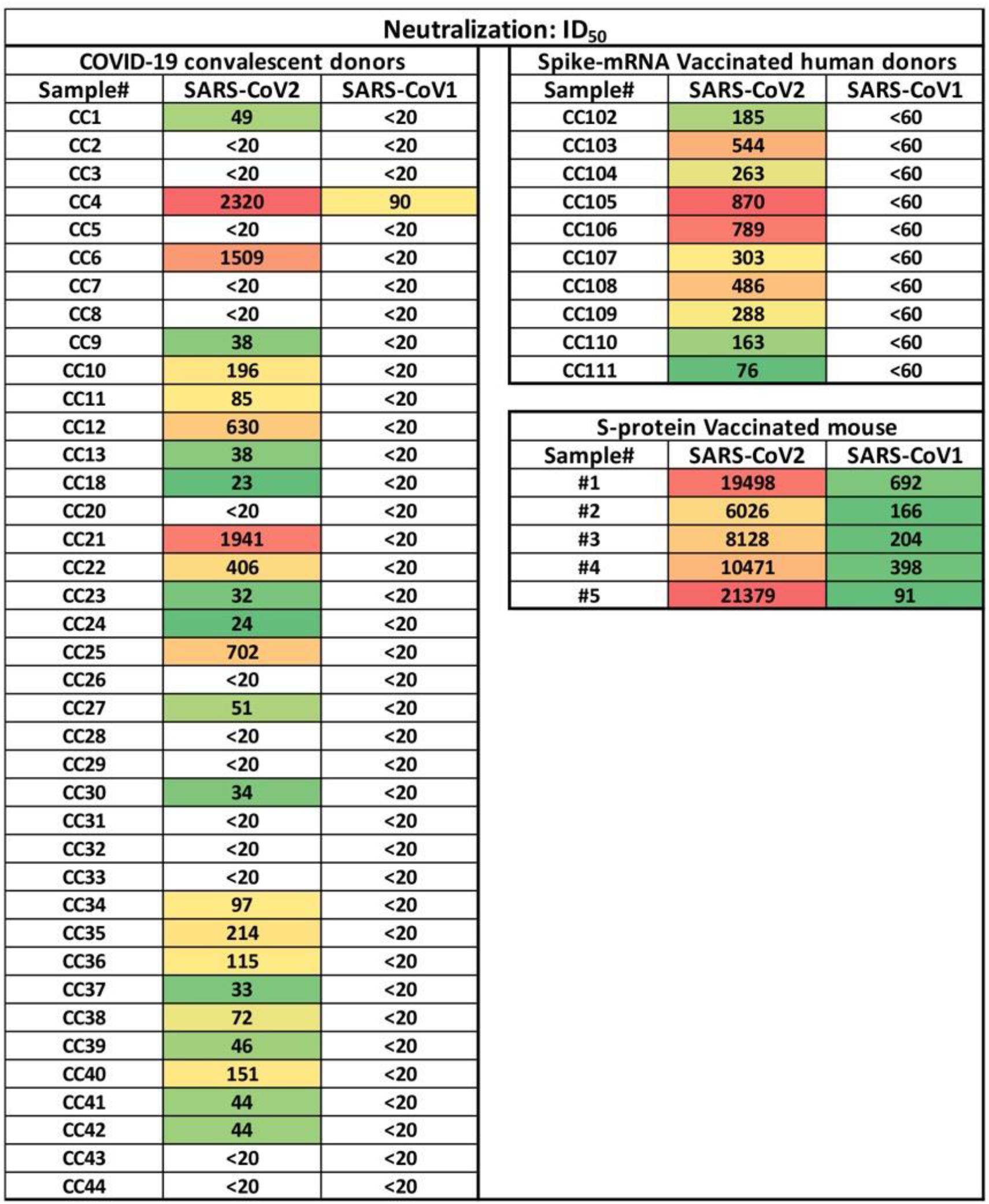
Neutralization of SARS-CoV-1 and SARS-CoV-2 by various immune sera. Neutralizing titer (ID_50_) of serum antibodies from COVID-19 convalescent patients (n = 39), Spike-mRNA vaccinated human donors (n = 10), and S-protein immunized mice (n = 5). Neutralization was tested against SARS-CoV-2 and SARS-CoV-1.

**Fig. S4.**
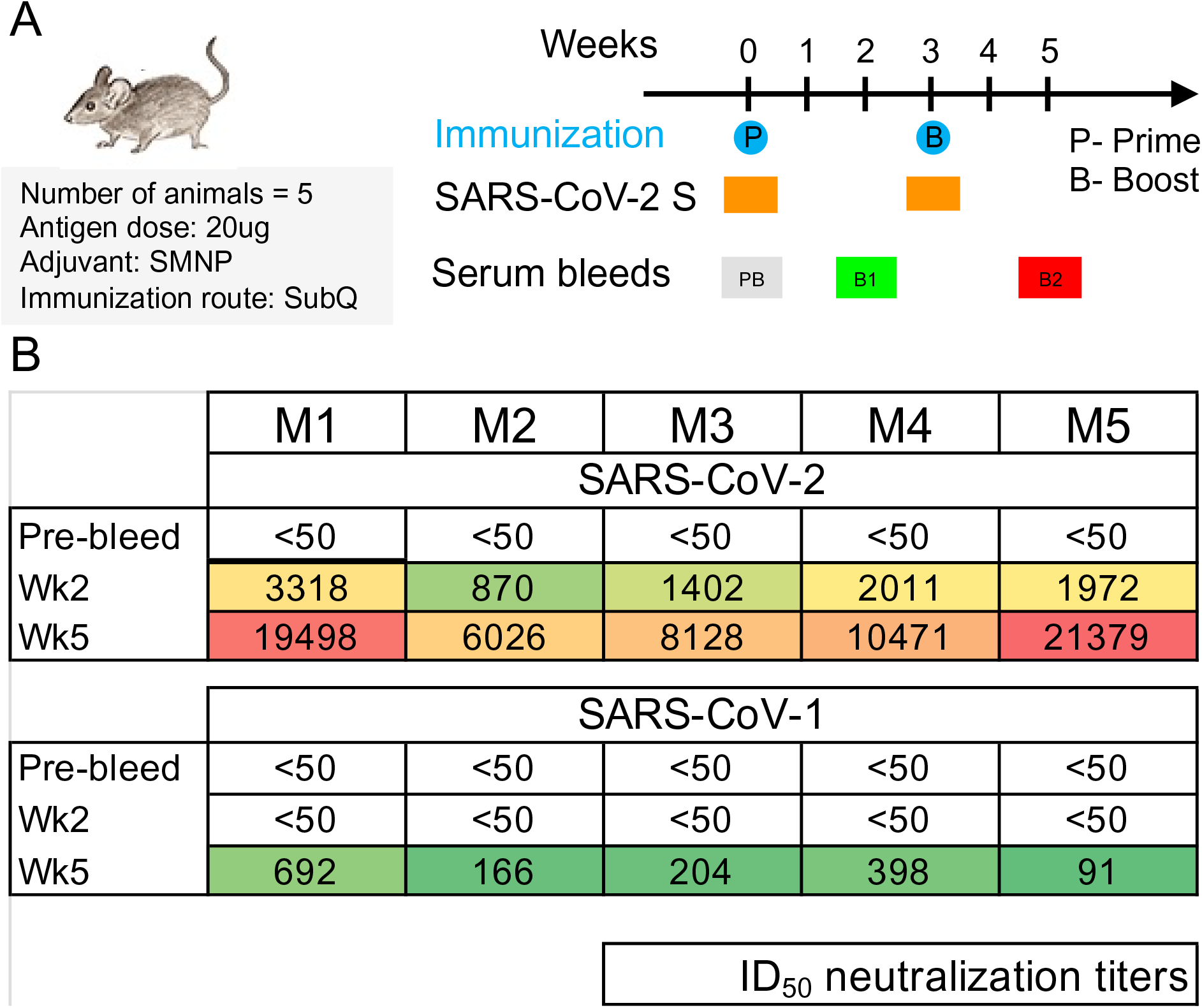
SARS-CoV-2 S-protein immunization in WT B6 mice and induction of neutralizing antibody responses against SARS-CoV-2 and SARS-CoV-1. **(A)** Immunization of B6 mice with SARS-CoV-2 S protein at week-0 (P-prime) and week-3 (B-boost). Groups of 5 mice were subcutaneously immunized twice (wk-0 prime and wk-3 boost) with 20ug of SARS-CoV-2 S protein along with SMNP (Isco-MPLA) adjuvant. **(B)** Neutralization of SARS-CoV-2 and SARS-CoV-1 by pre-bleed (PB), post-prime (B1-wk2) and post-boost (B2-wk5) immune sera. Serial dilutions of sera (starting dilution 1:50) were tested against SARS-CoV-2 and SARS-CoV-1 pseudotyped viruses in an ACE2 expressing cell-based assay and ID_50_ neutralization titers are shown.

**Fig. S5.**
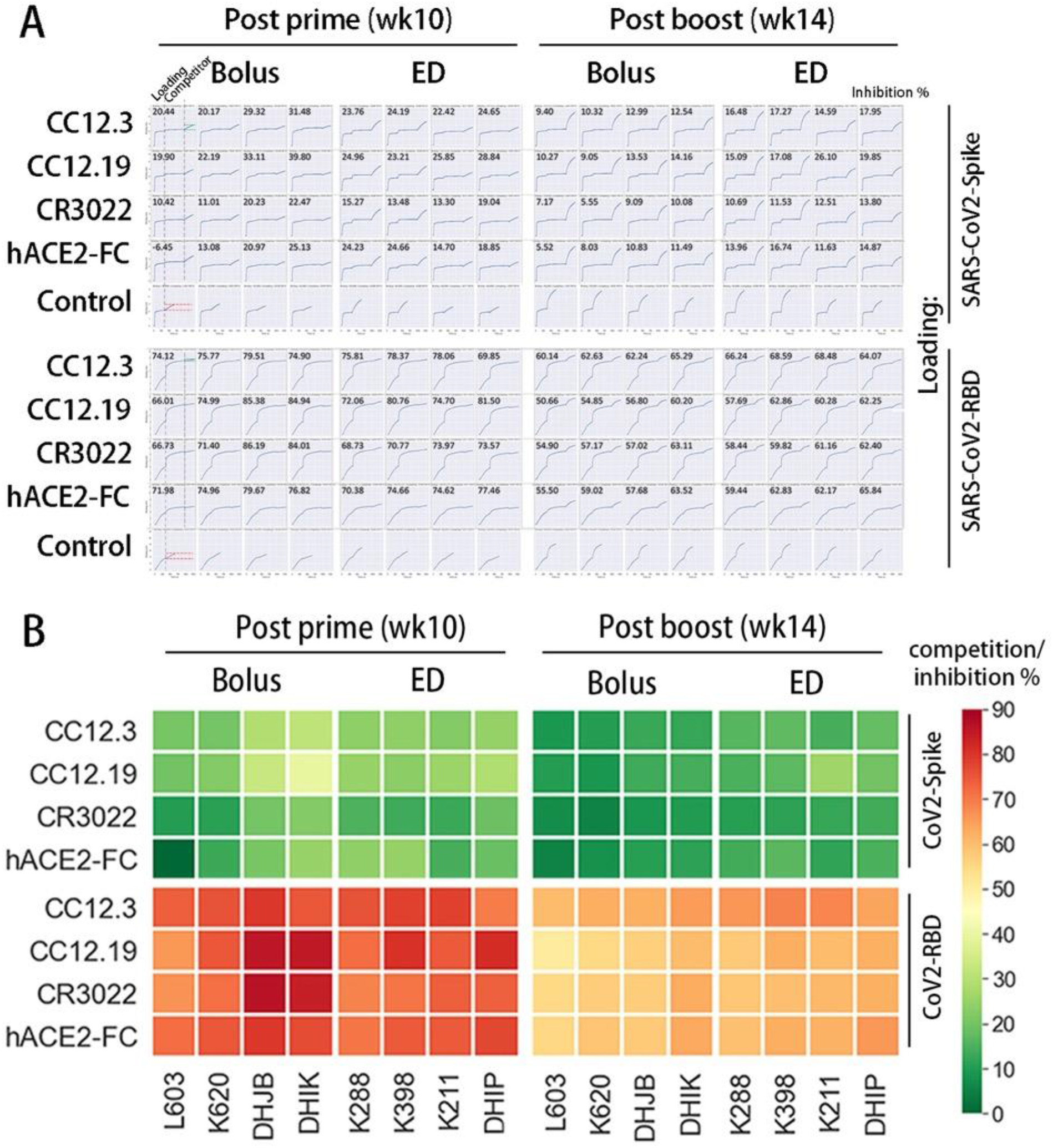
BLI binding competition of rhesus macaque immune sera with SARS-CoV-2 human nAbs and hACE2. **(A)** The sera from the SARS-CoV-2 S protein-immunized rhesus macaques post-prime (week10) and post-boost (week14) were evaluated for epitope competition with human SARS-CoV-2-specific mAbs and recombinant human hACE2 protein using Bio-layer Interferometry (BLI). Biotinylated SARS-CoV-2 spike (top rows) or RBD (bottom rows) proteins were captured using SA biosensors and incubated with the indicated mAbs at a saturating concentration of 100ug/mL for 10 mins followed by an incubation for 5 min in 1:50 diluted serum from SARS-CoV-2 S protein-immunized rhesus macaques. As controls, the biotinylated SARS-CoV-2 S or RBD proteins were captured using SA biosensors and incubated only in 1:50 diluted serum from S protein-immunized rhesus macaques. BLI raw traces are shown. The percent (%) binding inhibition is calculated with the formula: [percent (%) of inhibition in the BLI response = 1- (sera binding response in presence of the competitor antibody, indicated by green dash line / response of the corresponding control serum antibodies without the competitor antibody, indicated by red dash line)]**. (B)** Heatmap showing BLI competition-based epitope binning of NHP immune sera with human RBD-specific nAbs CC12.3, CC12.19, and CR3022, respectively, as well as with soluble hACE2-Fc. The competition experiments were performed with full-length SARS-CoV-2 S protein (CoV-2-Spike) and the monomeric SARS-CoV-2 RBD (CoV-2-RBD).

**Fig. S6.**
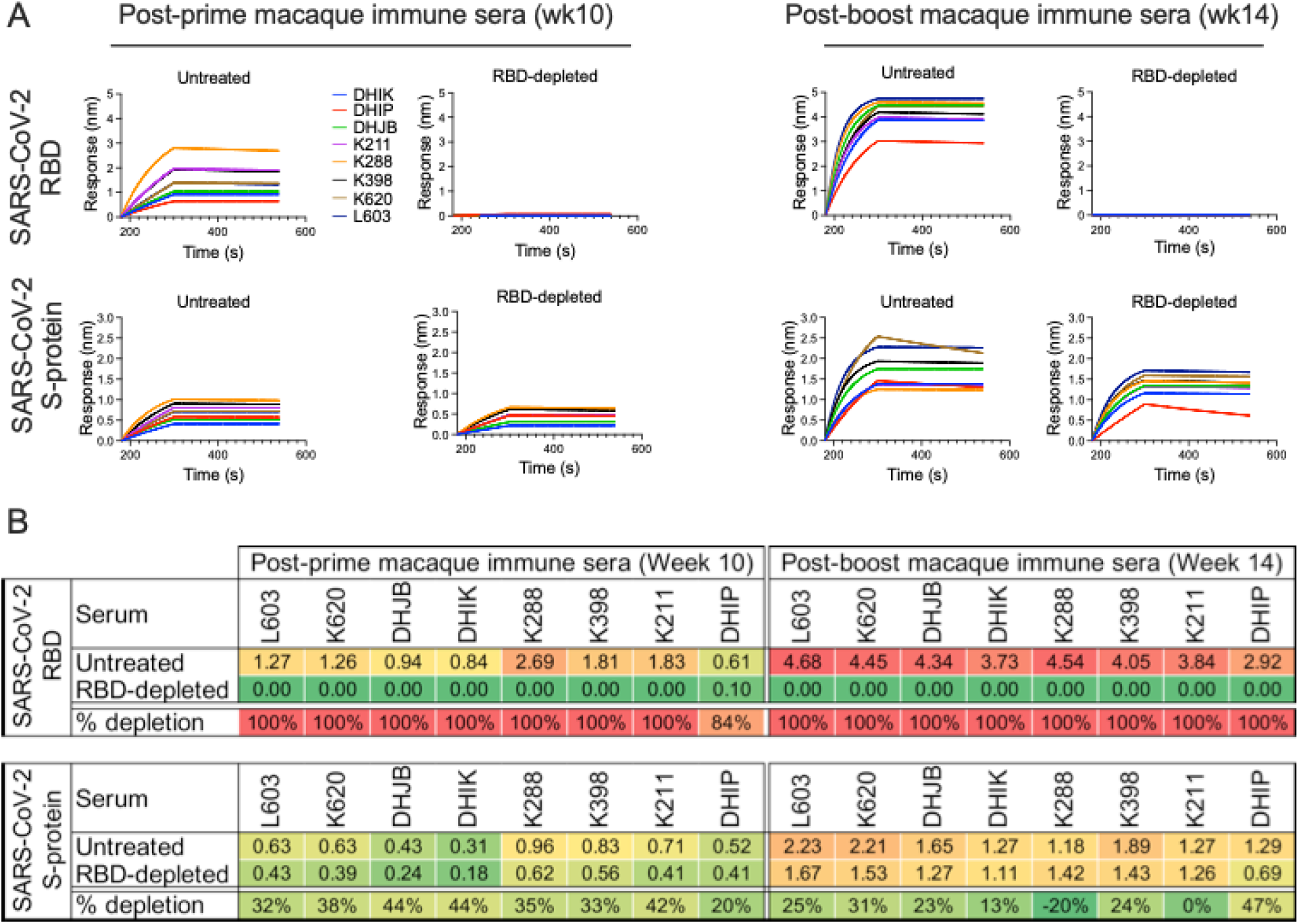
Depletion of RBD-specific antibodies in post-prime and post-boost macaque immune sera confirmed by BLI binding to monomeric SARS-CoV-2 RBD and SARS-CoV-2 S-protein. **(A)** BLI binding curves of post-prime (wk 10) and post-boost (wk 14) macaque immune sera with monomeric SARS-CoV-2 RBD and SARS-CoV-2 S-protein. Binding of both untreated and anti-RBD antibody adsorbed (RBD-depleted) immune sera are shown. The BLI revealed complete loss of binding to monomeric RBD in the anti-RBD depleted sera in both post-prime and post-boost immune sera. **(B)** Summary table showing the BLI binding responses and percent (%) depletion of anti-RBD antibodies in post-prime and post-boost immune sera upon adsorption of polyclonal antibodies with monomeric RBD. The BLI binding responses of the untreated and anti-RBD antibody adsorbed (RBD-depleted) immune sera with monomeric SARS-CoV-2 RBD and SARS-CoV-2 S-protein are shown. The % depletion of anti-RBD antibodies for each immune serum is calculated with the formula: [Percent (%) depletion = 1- (BLI binding response of the RBD-depleted immune sera / BLI binding response of the untreated immune sera) X 100]

**Fig. S7.**
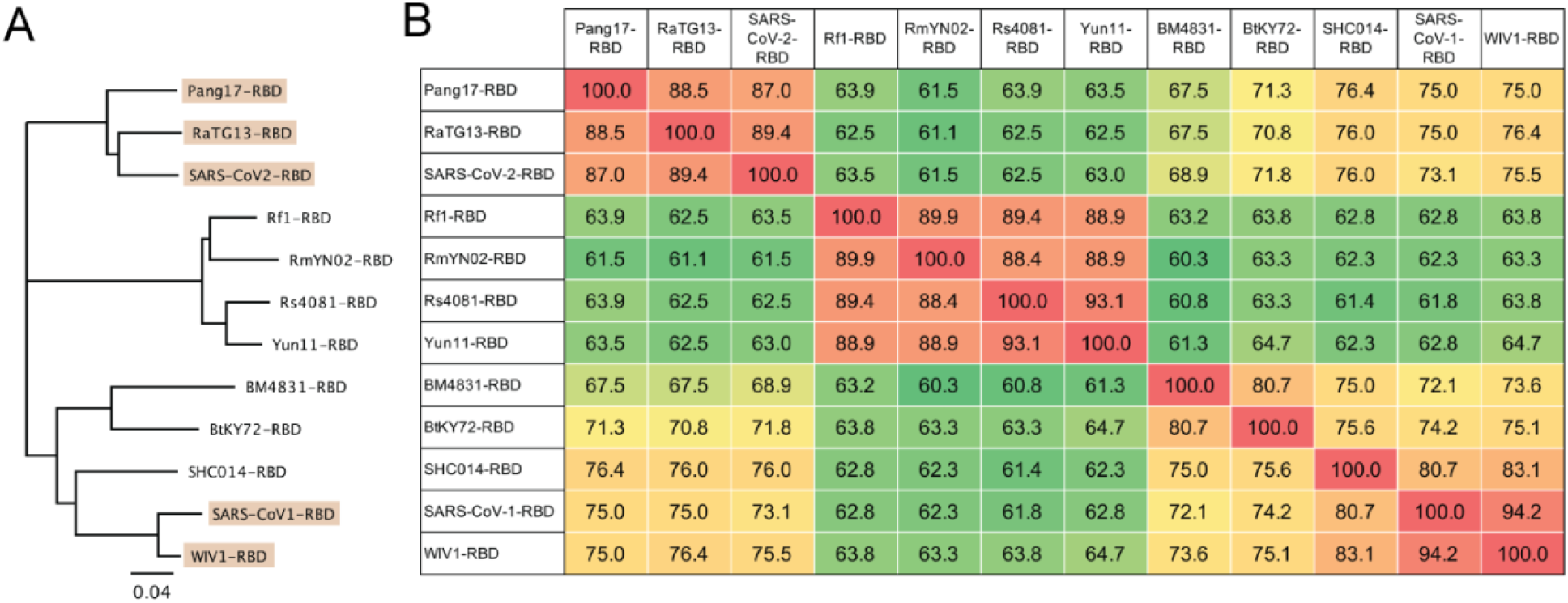
Phylogenetic relatedness of receptor binding domains of major sarbecovirus lineages. **(A)** Phylogenetic tree derived from amino-acid sequences of RBD regions of spikes of 12 representative sarbecovirus lineages. The five taxa highlighted in beige are SARS-like viruses that utilize ACE2 receptor for infection of the host cell and thus tested in neutralization assays. **(B)** Percent (%) identify matrix based on amino-acid sequences of RBD show relatedness among the sarbecovirus RBDs. Among the viruses tested for neutralization, SARS-CoV-2 RBD is closer to RatG13 (89.4%) and Pang17 (87%) RBDs compared to SARS-CoV-1 (73.1%) and WIV1 (75.5%) RBDs. WIV1 RBD is closer to SARS-CoV-1 (94.2%).

**Fig. S8.**
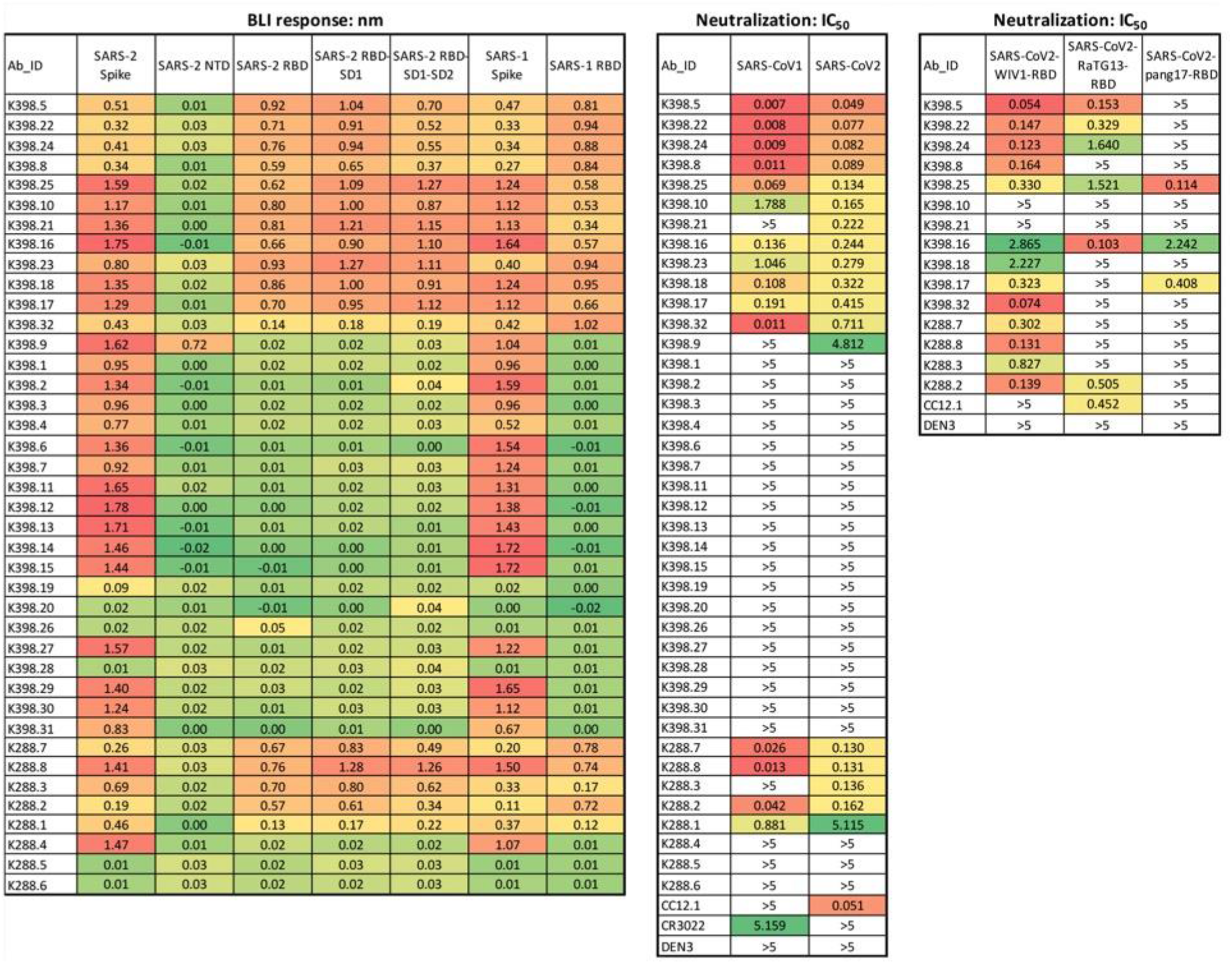
Binding and neutralization of vaccine-elicited monoclonal antibodies. Binding of mAbs isolated from immunized RM B cells toward S-proteins. BLI binding response of isolated mAbs from K398 and K288 against SARS-CoV-2 spike, SARS-CoV-2-NTD, RBD, RBD-SD1, RBD-SD1-SD2, SARS-CoV-1 spike, SARS-CoV1-RBD are shown in the left chart. The IC_50_ neutralization titers against SARS-CoV-1 and SARS-CoV-2 pseudoviruses are shown in the right chart. CC12.1, CR3022, and DEN3 were used as controls for the neutralization IC_50_ assays. The IC_50_ neutralization titers are shown for select cross-neutralizing mAbs with WIV1, RatG13 and pang17 RBD-SARS-CoV-2 chimeric pseudoviruses.

**Fig. S9.**
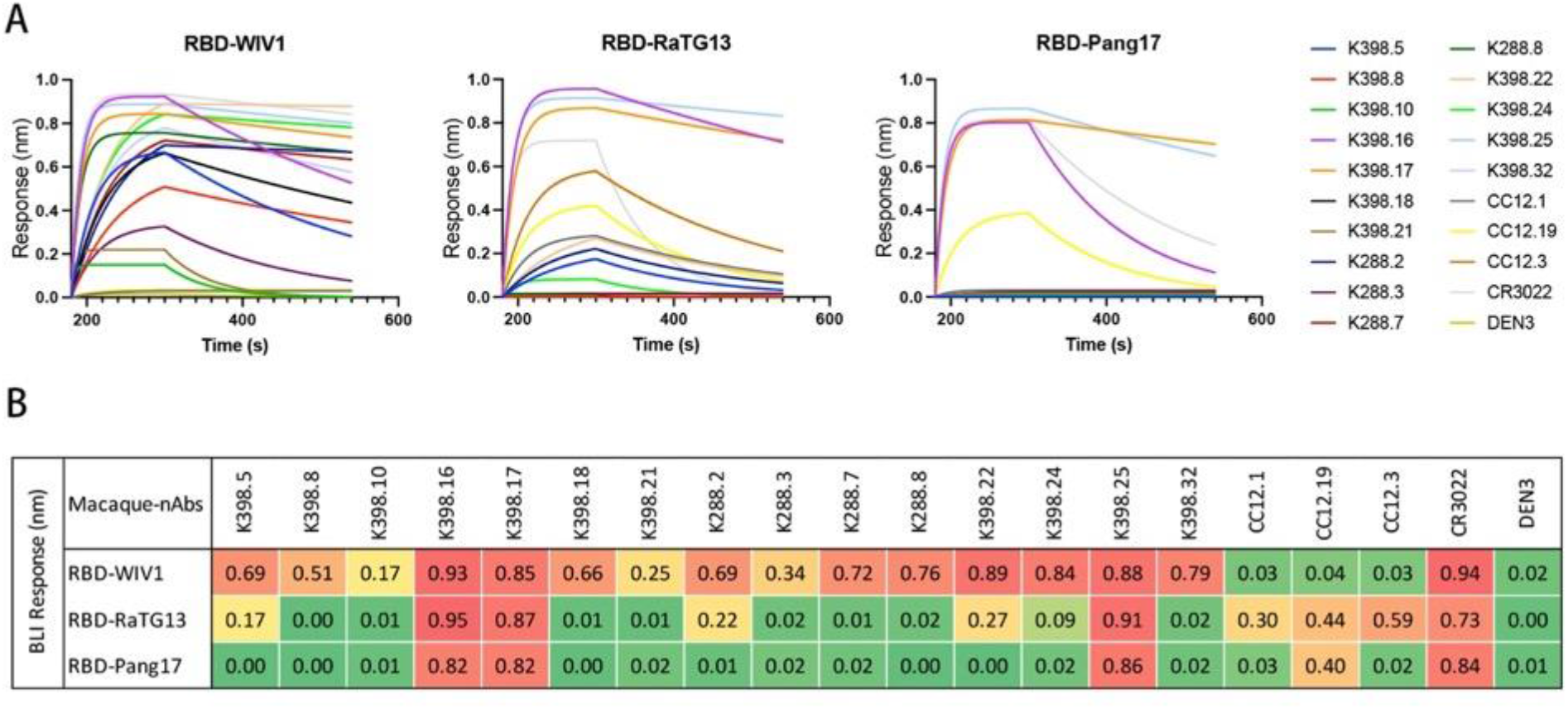
BLI binding of cross-reactive macaque antibodies with monomeric RBDs of ACE2- utilizing sarbecoviruses. **(A)** BLI binding curves of cross-reactive macaque antibodies with RBDs of WIV1, RatG13 and Pang17 sarbecoviruses. Human SARS-CoV-2 (CC12.1, CC12.19 and CC12.3), SARS-CoV-1 (CR3022) and unrelated Dengue (DEN3) specific antibodies were controls for the binding assay. **(B)** Summary table showing peak of the BLI binding responses for each antibody with given monomeric RBD proteins. The BLI binding of the macaque antibodies is consistent with their cross-neutralizing activities with the corresponding sarbecoviruses.

**Fig. S10.**
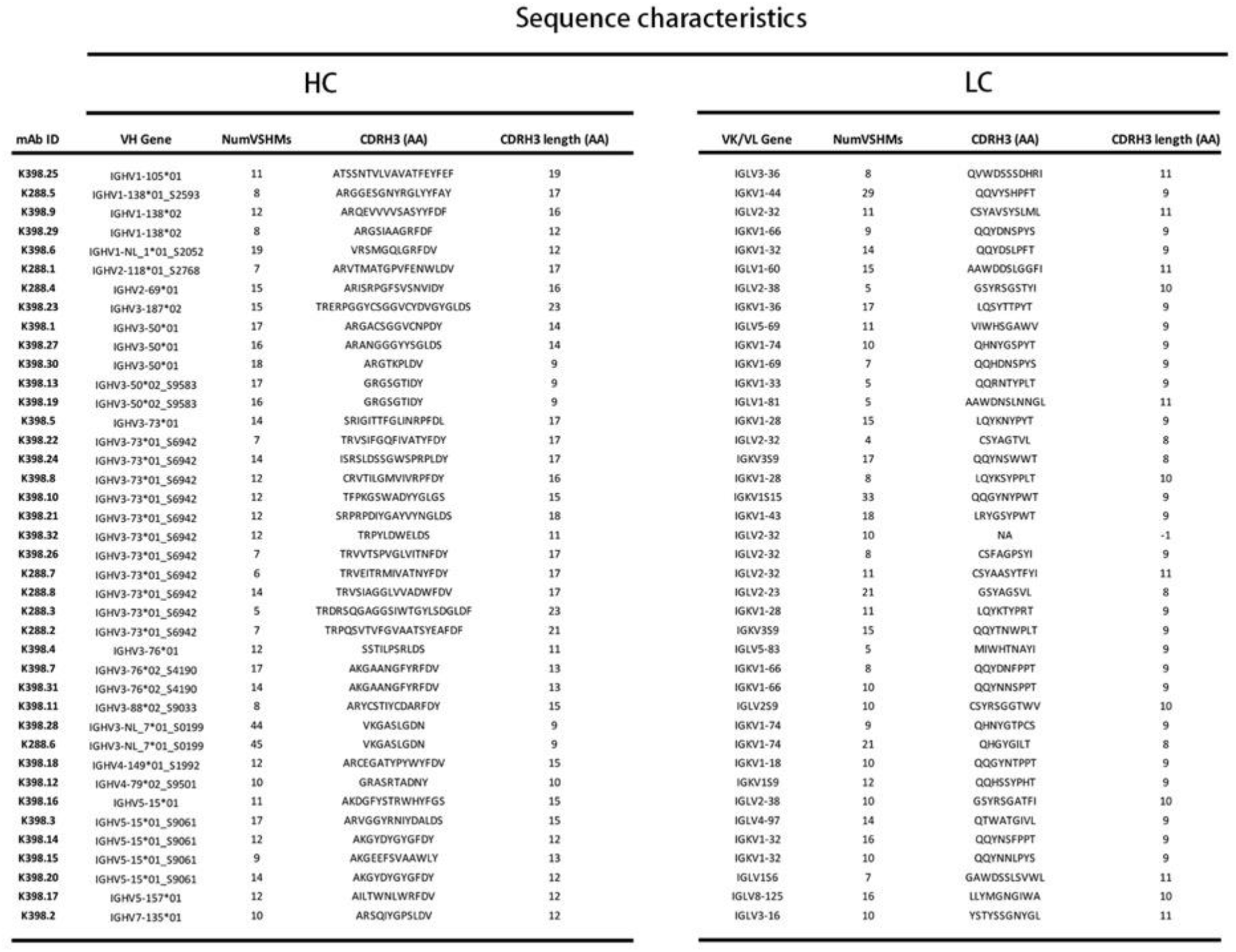
Immunogenetic properties of mAbs isolated from two SARS-CoV-2 S-protein immunized rhesus macaques, K398 and K288. The heavy (VH) and light (VK/VL) gene usage, somatic hypermutation levels, CDR3 sequences and lengths are shown for each mAb. The rhesus macaque (*Macaca mulatta*) germline database from (*62*) was used for all gene assignments.

**Fig. S11.**
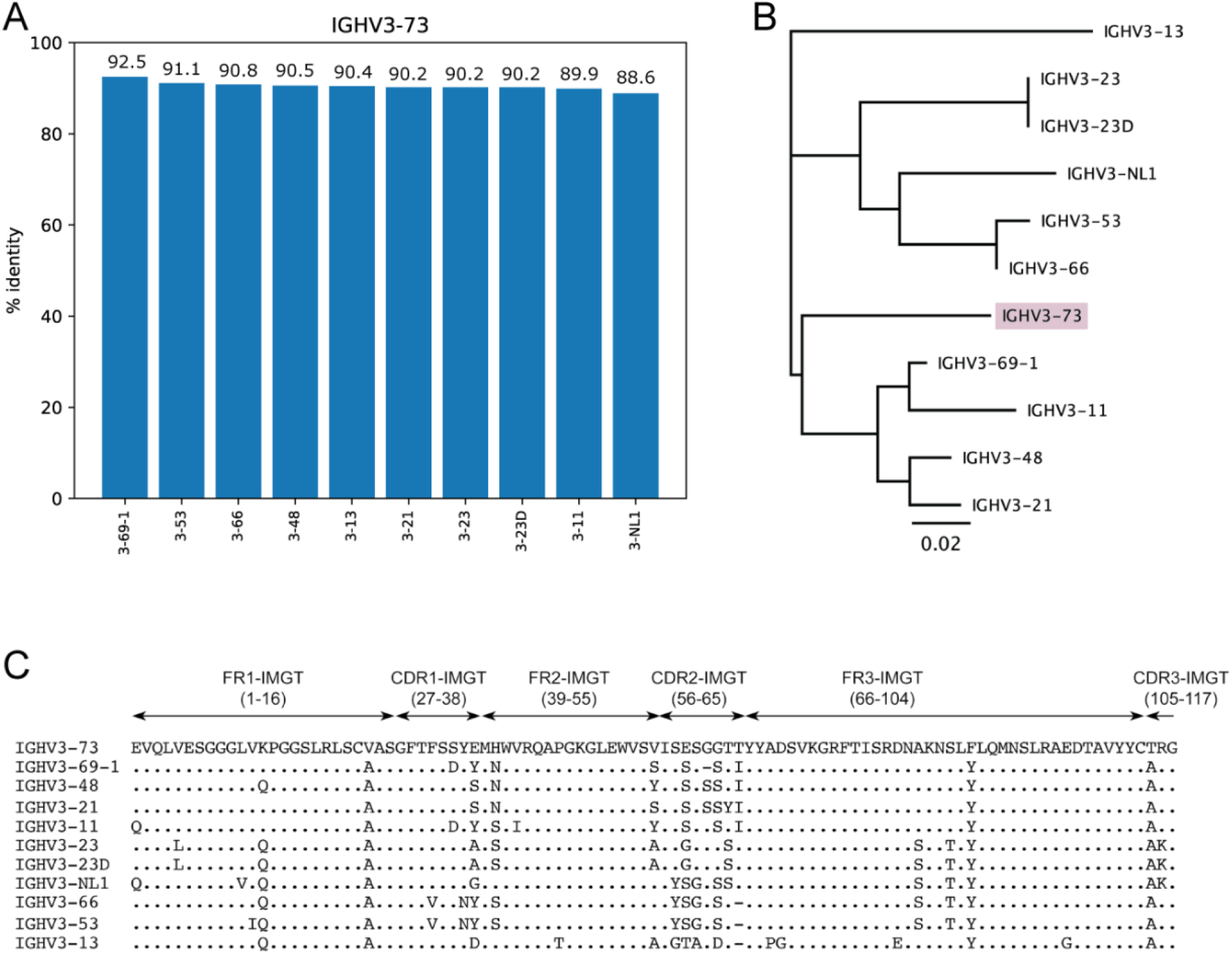
Sequence similarity of macaque IGHV3-73 germline with the closest human IGHV germline genes. **(A)** The nucleotide sequence of the macaque IGHV3-73 germline gene (utilized by macaque sarbecovirus cross-neutralizing antibodies) was compared with the entire human IGHV germline gene database and the bar plot shows percent (%) identity with the 10 closest human germline genes (3-69-1, 3-53, 3-66, 3-48, 3-13, 3-21, 3-23, 3-23D, 3-11 and 3-NL1). The % identity for each human IGHV germline gene with macaque IGHV3-73 germline is indicated. IGHV3-69-1 is a pseudogene and is not expressed. Notably, the macaque IGHV3-73 germline gene showed closest homology to the human IGHV3-53 and IGHV3-66 germline genes that are frequently enriched in human SARS-CoV-2 neutralizing antibodies. **(B)** Phylogenetic relatedness of macaque IGHV3-73 germline gene to the 10 closest human germline genes. The amino-acid sequences of the macaque IGHV3-73 germline gene and the closest human germline genes were used to generate the phylogenetic tree. **(C)** Amino-acid sequence alignment of macaque IGHV3- 73 germline V-gene with the 10 closest human germline genes. The human germline genes are arranged in the order of their relatedness (amino acid level) to the macaque IGHV3-73 germline gene.

**Fig. S12.**
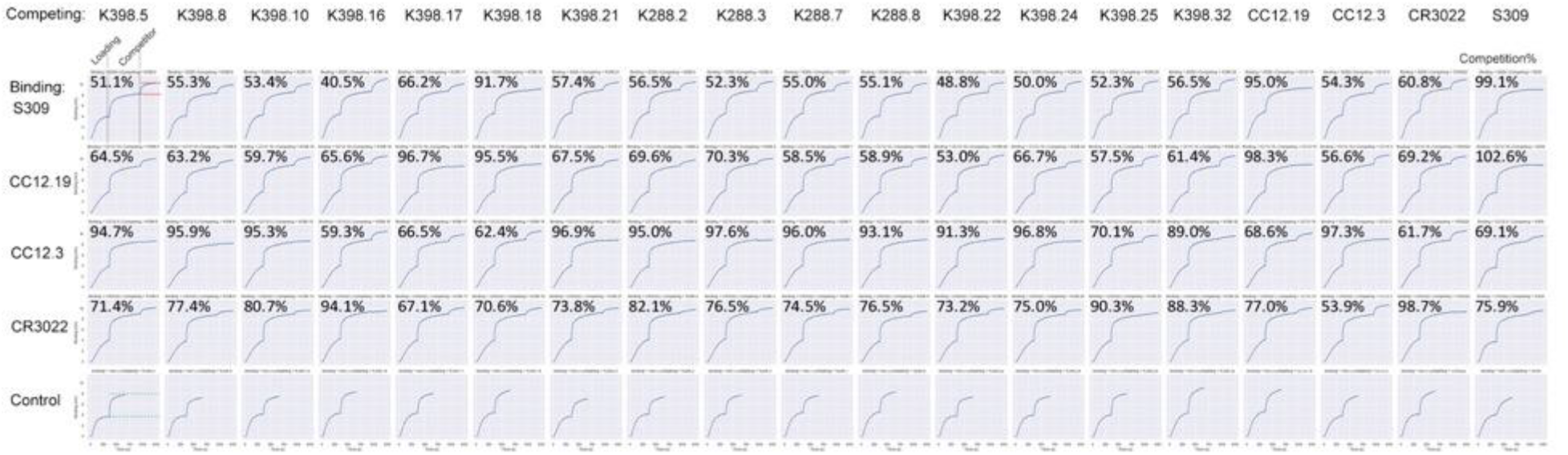
Epitope mapping of macaque cross-neutralizing mAbs by BLI competition binning. The nAbs isolated from the SARS-CoV-2 S protein immunized rhesus macaques were evaluated for epitope competition with human SARS-CoV-2 specific mAbs, CC12.19, CC12.3 as well as cross-reactive S309 and CR3022 mAbs. His-tagged SARS-CoV-2 RBD protein was captured using anti-His biosensors and incubated with the indicated mAbs at a saturating concentration of 100ug/mL for 10 mins and followed by an incubation for 5 min in the nAbs at concentration of 25ug/mL. BLI traces are shown for each binding. The binding inhibition % is calculated with the formula: percent (%) of inhibition in the BLI binding response = 1- (response in presence of the competitor antibody, indicated by green dashed line / response of the corresponding control antibody without the competitor antibody, indicated by red dashed line).

**Fig. S13.**
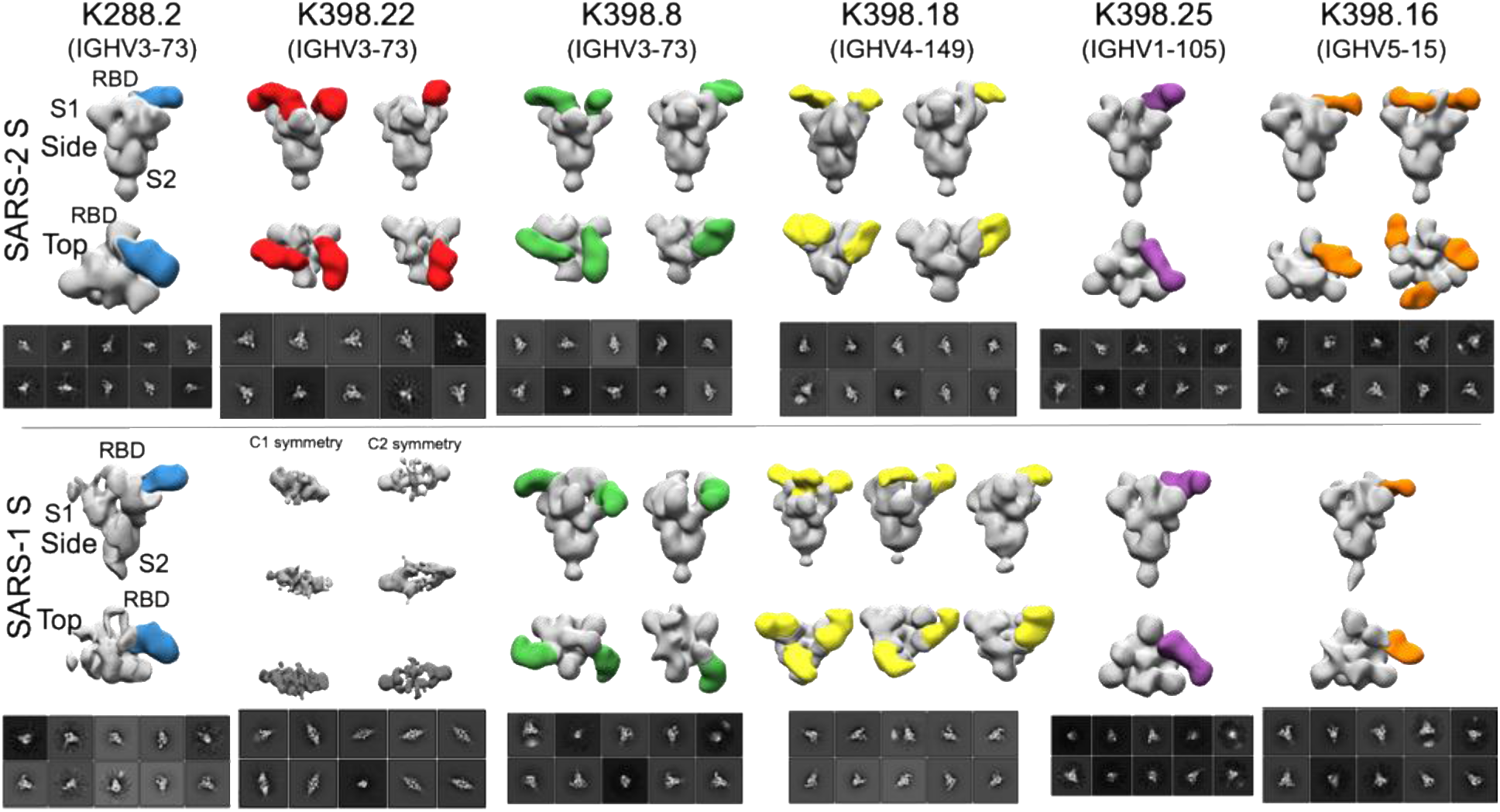
Epitope mapping of sarbecovirus cross-neutralizing macaque mAbs bound to SARS-CoV-2 and SARS-CoV-1 S-proteins by negative stain electron microscopy. Electron microscopy (EM) 3D reconstructions of sarbecovirus cross-neutralizing macaque antibodies K288.2 (blue), K398.22 (red), K398.8 (green), K398.18 (yellow), K398.25 (purple) and K398.16 (orange) in complex with SARS-CoV-2 (upper panel) and SARS-CoV-1 (lower panel) S-proteins. The IGHV gene usage for each macaque cross-nAb is indicated. The EM 3D reconstructions of Fab and S-protein complexes were generated from nsEM 2D class average images shown for each mAb, under 3D reconstruction models. The spike S1 and S2 subunits and RBD are labelled for representative cross-nAb, K288.2.

**Fig. S14.**
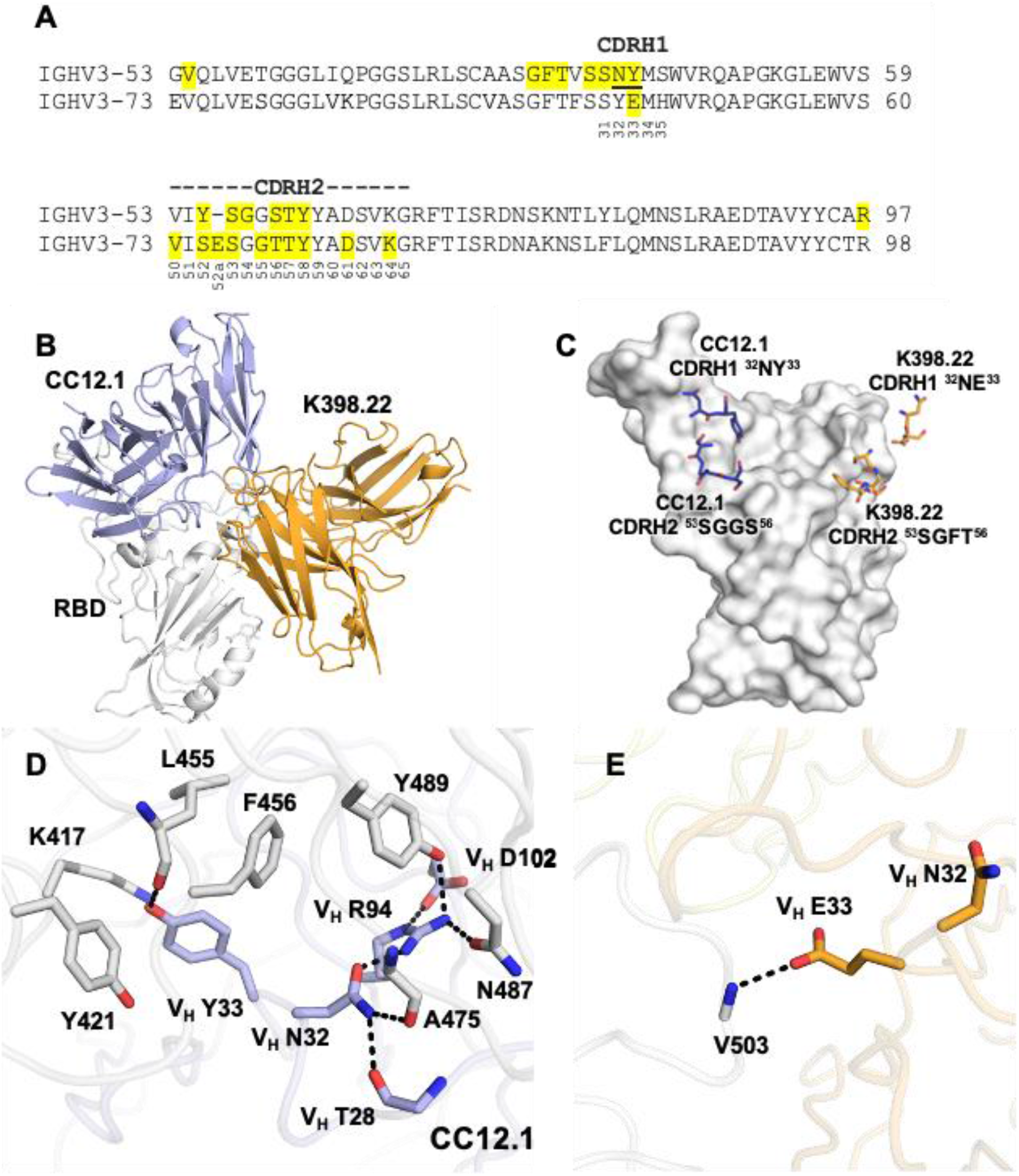
Macaque IGHV3-73 is closest to human IGHV3-53 in sequence but differs in the critical RBD-interacting human germline ‘NY’ motif and adopts a completely different binding mode. **(A)** Sequence alignment between the germlines (human IGHV3-53 and macaque IGHV3-73) that encode shared antibody responses. Paratope residues of IGHV3-53 antibodies targeting SARS-CoV-2 in the canonical mode (*22*) as well as macaque IGHV3-73 antibodies in this study are highlighted in yellow. Although the two germline sequences are similar (∼90% sequence identity), the critical RBD-interacting human IGHV3-53 ‘NY’ motif (underlined) is replaced by ‘YE’ in macaque IGHV3-73 and does not bind in the same mode. **(B)** Macaque IGHV3-73 and human IGHV3-53 antibodies adopt very different binding modes. K398.22 is used to represent the macaque IGHV3-73 antibodies and CC12.1 for human IGHV3-53 antibodies (PDB 6XC2). **(C-E)** Detailed interactions between SARS-CoV-2 RBD (white) and the CDRH1 ‘NY’ motif of a human IGHV3-53 antibody CC12.1 (PDB 6XC2) and CDRH1 ‘NE’ motif of macaque IGHV3-73 antibody K398.22. Interactions of corresponding (D) CC12.1 and (E) K398.22 CDRH1 residues with RBD are indicated. Hydrogen bonds and salt bridges are represented by black dashed lines.

**Fig. S15.**
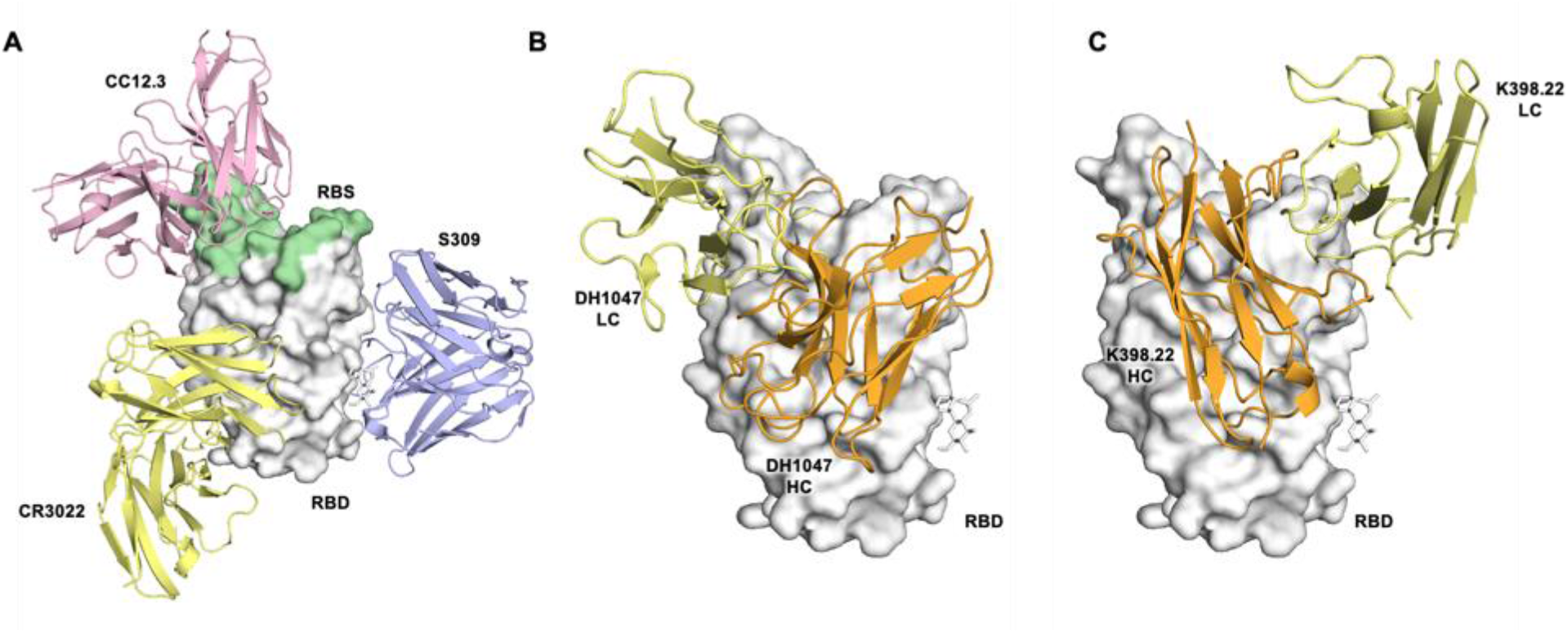
Distinct binding modes of human and macaque RBD directed SARS-CoV-2 neutralizing antibodies. SARS-CoV-2 RBD is represented as a white surface. RBD molecules are in the same orientation in each panel. Antibodies are shown as cartoons in different colors. For clarity, only the variable domains of the antibodies are shown. **(A)** Binding modes of human SARS-CoV-2 neutralizing antibodies (CC12.3: pink; CR3022: yellow; S309: blue) with RBD. The N343 glycan is shown in stick representation. Each antibody interacts with the RBD in a different binding mode. The RBS on RBD is highlighted in green**. (B-C)** Comparison between the binding of (B) DH1047 and (C) K398.22. For both antibodies, the heavy chains are in orange and light chains in yellow. The heavy chains for the two antibodies target a similar region on the RBD but in a different binding orientation.

**Fig. S16.**
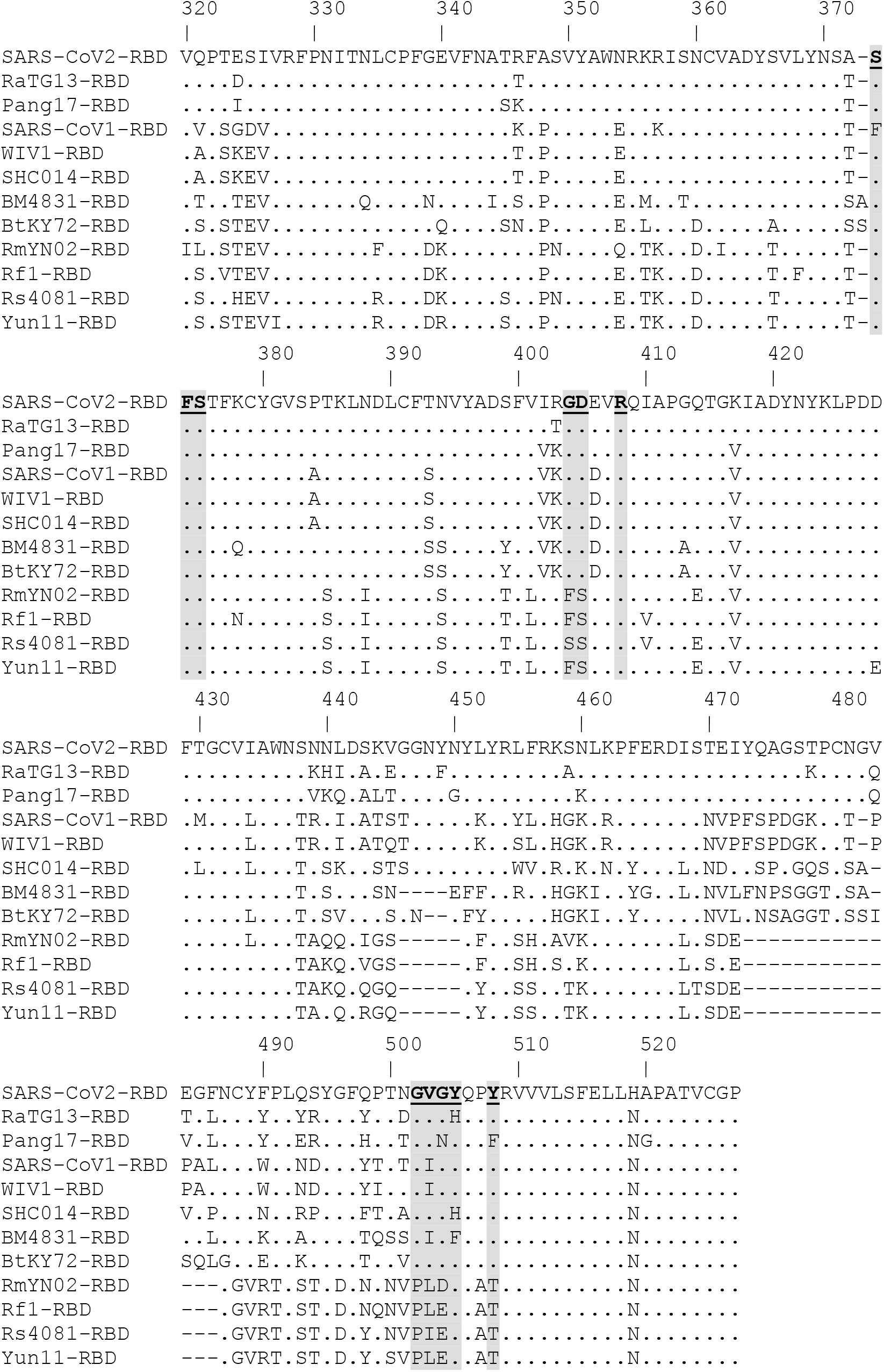
Conservation of the RBD bnAb epitope recognized by macaque IGHV3-73 encoded cross-nAbs. Amino-acid sequence alignment of sarbecovirus RBD regions from 12 major lineages, as indicated on the left. Sequence conservation of the epitope residues (highlighted in grey) in SARS-related strains recognized by macaque IGHV3-73 encoded bnAbs.

## REFERENCES AND NOTES

1. L. Dai, G. F. Gao, Viral targets for vaccines against COVID-19. Nat Rev Immunol 21, 73–82 (2020).

2. F. Amanat, F. Krammer, SARS-CoV-2 Vaccines: Status Report. Immunity 52, 583–589 (2020).

3. F. Krammer, SARS-CoV-2 vaccines in development. Nature 586, 516–527 (2020).

4. S. F. Ahmed, A. A. Quadeer, M. R. McKay, Preliminary identification of potential vaccine targets for the COVID-19 coronavirus (SARS-CoV-2) based on SARS-CoV immunological studies. Viruses 12, 254 (2020).

5. P. J. Klasse, D. F. Nixon, J. P. Moore, Immunogenicity of clinically relevant SARS-CoV-2 vaccines in nonhuman primates and humans. Sci Adv 7, eabe8065 (2021).

6. L. A. Jackson et al., An mRNA vaccine against SARS-CoV-2 - Preliminary report. N Engl J Med 383, 1920–1931 (2020).

7. F. P. Polack et al., Safety and efficacy of the BNT162b2 mRNA Covid-19 vaccine. N Engl J Med 383, 2603–2615 (2020).

8. Y. Cao et al., Humoral immune response to circulating SARS-CoV-2 variants elicited by inactivated and RBD-subunit vaccines. Cell Res 7, 732–741 (2021).

9. Z. Wang et al., mRNA vaccine-elicited antibodies to SARS-CoV-2 and circulating variants. Nature 592, 616–622 (2021).

10. D. F. Robbiani et al., Convergent antibody responses to SARS-CoV-2 in convalescent individuals. Nature 584, 437–442 (2020).

11. T. F. Rogers et al., Isolation of potent SARS-CoV-2 neutralizing antibodies and protection from disease in a small animal model. Science 369, 956–963 (2020).

12. A. Z. Wec et al., Broad neutralization of SARS-related viruses by human monoclonal antibodies. Science 369, 731–736 (2020).

13. B. Ju et al., Human neutralizing antibodies elicited by SARS-CoV-2 infection. Nature 584, 115–119 (2020).

14. R. Shi et al., A human neutralizing antibody targets the receptor-binding site of SARS- CoV-2. Nature 584, 120–124 (2020).

15. S. Du et al., Structurally resolved SARS-CoV-2 antibody shows high efficacy in severely infected hamsters and provides a potent cocktail pairing strategy. Cell 183, 1013–1023 e1013 (2020).

16. S. J. Zost et al., Potently neutralizing and protective human antibodies against SARS- CoV-2. Nature 584, 443–449 (2020).

17. Y. Wu et al., A noncompeting pair of human neutralizing antibodies block COVID-19 virus binding to its receptor ACE2. Science 368, 1274–1278 (2020).

18. P. J. M. Brouwer et al., Potent neutralizing antibodies from COVID-19 patients define multiple targets of vulnerability. Science 369, 643–650 (2020).

19. J. Hansen et al., Studies in humanized mice and convalescent humans yield a SARS- CoV-2 antibody cocktail. Science 369, 1010–1014 (2020).

20. C. O. Barnes et al., Structures of Human Antibodies Bound to SARS-CoV-2 Spike Reveal Common Epitopes and Recurrent Features of Antibodies. Cell 182, 828–842 e816 (2020).

21. Y. Cao et al., Potent neutralizing antibodies against SARS-CoV-2 identified by high- throughput single-cell sequencing of convalescent patients’ B cells. Cell 182, 73–84 e16 (2020).

22. M. Yuan et al., Structural basis of a shared antibody response to SARS-CoV-2. Science 369, 1119–1123 (2020).

23. M. Yuan et al., Structural and functional ramifications of antigenic drift in recent SARS- CoV-2 variants. bioRxiv, (2021).

24. C. O. Barnes et al., SARS-CoV-2 neutralizing antibody structures inform therapeutic strategies. Nature 588, 682–687 (2020).

25. . S. I. Kim et al., Stereotypic neutralizing VH antibodies against SARS-CoV-2 spike protein receptor binding domain in patients with COVID-19 and healthy individuals. Sci. Tranl. Med. 13, eabd6990 (2021).

26. E. Andreano et al., SARS-CoV-2 escape *in vitro* from a highly neutralizing COVID-19 convalescent plasma. bioRxiv, 2020.2012.2028.424451 (2020).

27. Y. Weisblum et al., Escape from neutralizing antibodies by SARS-CoV-2 spike protein variants. Elife 9, e61312 (2020).

28. C. K. Wibmer et al., SARS-CoV-2 501Y.V2 escapes neutralization by South African COVID-19 donor plasma. bioRxiv, (2021).

29. J. R. Mascola, B. S. Graham, A. S. Fauci, SARS-CoV-2 Viral Variants-Tackling a Moving Target. JAMA, 325(**13**) 1261–1262 (2021).

30. P. Wang et al., Antibody Resistance of SARS-CoV-2 Variants B.1.351 and B.1.1.7. Nature 593,130–135 (2021).

31. K. M. Cirelli et al., Slow delivery immunization enhances HIV neutralizing antibody and germinal center responses via modulation of immunodominance. Cell 177, 1153–1171 e1128 (2019).

32. T. J. Moyer et al., Engineered immunogen binding to alum adjuvant enhances humoral immunity. Nat. Med. 26, 430–440 (2020).

33. H. H. Tam et al., Sustained antigen availability during germinal center initiation enhances antibody responses to vaccination. Proc. Natl. Acad. Sci. U.S.A.113, E6639–E6648 (2016).

34. D. E. Anderson et al., Lack of cross-neutralization by SARS patient sera towards SARS- CoV-2. Emerg. Microb. Infect. 9, 900–902 (2020).

35. H. Lv et al., Cross-reactive antibody response between SARS-CoV-2 and SARS-CoV infections. bioRxiv, (2020).

36. G. Song et al., Cross-reactive serum and memory B-cell responses to spike protein in SARS-CoV-2 and endemic coronavirus infection. Nat. Com. 12, 2938 (2021).

37. L. Stamatatos et al., mRNA vaccination boosts cross-variant neutralizing antibodies elicited by SARS-CoV-2 infection. Science 372, 1413–1418 (2021).

38. K. Worzner et al., Adjuvanted SARS-CoV-2 spike protein elicits neutralizing antibodies and CD4 T cell responses after a single immunization in mice. EBioMedicine 63, 103197 (2021).

39. J. Tong, C. Zhu, H. Lai, C. Feng, D. Zhou, Potent Neutralization Antibodies Induced by a Recombinant Trimeric Spike Protein Vaccine Candidate Containing PIKA Adjuvant for COVID-19. Vaccines (Basel) 9, 296 (2021).

40. A. E. Powell et al., A single immunization with spike-functionalized ferritin vaccines elicits neutralizing antibody responses against SARS-CoV-2 in mice. ACS Cent Sci 7, 183–199 (2021).

41. J. ter Meulen et al., Human monoclonal antibody combination against SARS coronavirus: synergy and coverage of escape mutants. PLoS Med 3, e237 (2006).

42. D. Angeletti et al., Defining B cell immunodominance to viruses. Nat. Immunol. 18, 456–463 (2017).

43. H. L. Turner et al., Disassembly of HIV envelope glycoprotein trimer immunogens is driven by antibodies elicited via immunization. bioRxiv, 2021.2002.2016.431310 (2021).

44. K. O. Saunders et al., Neutralizing antibody vaccine for pandemic and pre-emergent coronaviruses. Nature 594, 553–559, (2021).

45. M. Letko, A. Marzi, V. Munster, Functional assessment of cell entry and receptor usage for SARS-CoV-2 and other lineage B betacoronaviruses. Nat Microbiol 5, 562-569 (2020).

46. V. D. Menachery et al., A SARS-like cluster of circulating bat coronaviruses shows potential for human emergence. Nat. Med. 21, 1508–1513 (2015).

47. D. Pinto et al., Cross-neutralization of SARS-CoV-2 by a human monoclonal SARS-CoV antibody. Nature 583, 290–295 (2020).

48. A. A. Cohen et al., Mosaic nanoparticles elicit cross-reactive immune responses to zoonotic coronaviruses in mice. Science 371, 735–741 (2021).

49. P. S. Arunachalam et al., Adjuvanting a subunit COVID-19 vaccine to induce protective immunity. Nature 594, 253–258, (2021).

50. J. R. Francica et al., Vaccination with SARS-CoV-2 spike protein and AS03 adjuvant induces rapid anamnestic antibodies in the lung and protects against virus challenge in nonhuman primates. bioRxiv, (2021).

51. J. G. Liang et al., S-Trimer, a COVID-19 subunit vaccine candidate, induces protective immunity in nonhuman primates. Nat. Com. 12, 1346 (2021).

52. J. H. Tian, et al., SARS-CoV-2 spike glycoprotein vaccine candidate NVX-CoV2373 immunogenicity in baboons and protection in mice. Nat. Com. 12, 372 (2021).

53. M. G. Joyce et al., SARS-CoV-2 ferritin nanoparticle vaccines elicit broad SARS coronavirus immunogenicity. bioRxiv, (2021).

54. A. C. Walls et al., Elicitation of potent neutralizing antibody responses by designed protein nanoparticle vaccines for SARS-CoV-2. Cell 183, 1367–1382 e1317 (2020).

55. K. S. Corbett, et al., Evaluation of the mRNA-1273 Vaccine against SARS-CoV-2 in Nonhuman Primates. The N Engl J Med 383, 1544-1555 (2020).

56. K. S. Corbett, et al., Immune correlates of protection by mRNA-1273 immunization against SARS-CoV-2 infection in nonhuman primates. bioRxiv, (2021).

57. K. McMahan et al., Correlates of protection against SARS-CoV-2 in rhesus macaques. Nature 590, 630–634 (2021).

58. F. Krammer, Correlates of protection from SARS-CoV-2 infection. Lancet 397, 1421–1423 (2021).

59. A. Addetia et al., Neutralizing antibodies correlate with protection from SARS-CoV-2 in humans during a fishery vessel outbreak with a high attack rate. J Clin Microbiol 58, (2020).

60. V. J. Hall et al., SARS-CoV-2 infection rates of antibody-positive compared with antibody- negative health-care workers in England: a large, multicentre, prospective cohort study (SIREN). Lancet 397, 1459–1469 (2021).

61. B. Briney, A. Inderbitzin, C. Joyce, D. R. Burton, Commonality despite exceptional diversity in the baseline human antibody repertoire. Nature 566, 393–397 (2019).

62. N. Vazquez Bernat et al., Rhesus and cynomolgus macaque immunoglobulin heavy-chain genotyping yields comprehensive databases of germline VDJ alleles. Immunity 54, 355–366 e354 (2021).

63. J. G. Jardine et al., HIV-1 VACCINES. Priming a broadly neutralizing antibody response to HIV-1 using a germline-targeting immunogen. Science 349, 156-161 (2015).

64. K. M. Konrath et al., Nucleic acid delivery of immune-focused SARS-CoV-2 nanoparticles drive rapid and potent immunogenicity capable of single-dose protection. bioRxiv, 2021.2004.2028.441474 (2021).

65. S. W. de Taeye et al., Immunogenicity of Stabilized HIV-1 Envelope Trimers with Reduced Exposure of Non-neutralizing Epitopes. Cell 163, 1702–1715 (2015).

66. H. Duan et al., Glycan masking focuses immune responses to the HIV-1 CD4-binding site and enhances elicitation of VRC01-Class precursor antibodies. Immunity 49, 301–311 e305 (2018).

67. B. Briney et al., Tailored immunogens direct affinity maturation toward HIV neutralizing antibodies. Cell 166, 1459–1470 e1411 (2016).

68. D. Li et al., The functions of SARS-CoV-2 neutralizing and infection-enhancing antibodies in vitro and in mice and nonhuman primates. bioRxiv, (2021).

69. E. C. Lee et al., Complete humanization of the mouse immunoglobulin loci enables efficient therapeutic antibody discovery. Nat. Biotech. 32, 356–363 (2014).

70. M. J. Osborn et al., High-affinity IgG antibodies develop naturally in Ig-knockout rats carrying germline human IgH/Igkappa/Iglambda loci bearing the rat CH region. Journal of Immunol 190, 1481–1490 (2013).

71. A. J. Murphy et al., Mice with megabase humanization of their immunoglobulin genes generate antibodies as efficiently as normal mice. Proc. Natl. Acad. Sci. U.S.A. 111, 5153–5158 (2014).

72. M. Tian et al., Induction of HIV neutralizing antibody lineages in mice with diverse precursor repertoires. Cell 166, 1471–1484 e1418 (2016).

73. L. Verkoczy, F. W. Alt, M. Tian, Human Ig knockin mice to study the development and regulation of HIV-1 broadly neutralizing antibodies. Immunol. Rev. 275, 89–107 (2017).

74. J. Lan et al., Structure of the SARS-CoV-2 spike receptor-binding domain bound to the ACE2 receptor. Nature 581, 215–220 (2020).

## REFERENCES

75. P. Zhou et al., A protective broadly cross-reactive human antibody defines a conserved site of vulnerability on beta-coronavirus spikes. bioRxiv, (2021).

76. R. N. Kirchdoerfer et al., Pre-fusion structure of a human coronavirus spike protein. Nature 531, 118–121 (2016).

77. D. Wrapp et al., Cryo-EM structure of the 2019-nCoV spike in the prefusion conformation. Science 367, 1260–1263 (2020).

78. D. Sok et al., Recombinant HIV envelope trimer selects for quaternary-dependent antibodies targeting the trimer apex. Proc. Natl. Acad. Sci. U.S.A. 111, 17624–17629 (2014).

79. T. Tiller et al., Efficient generation of monoclonal antibodies from single human B cells by single cell RT-PCR and expression vector cloning. J Immunol. Met. 329, 112–124 (2008).

80. J. Gorman et al., Isolation and structure of an antibody that fully neutralizes isolate SIVmac239 reveals functional similarity of SIV and HIV glycan shields. Immunity 51, 724–734 e724 (2019).

81. C. Suloway et al., Automated molecular microscopy: the new Leginon system. J Struct Biol 151, 41–60 (2005).

82. N. R. Voss, C. K. Yoshioka, M. Radermacher, C. S. Potter, B. Carragher, DoG Picker and TiltPicker: software tools to facilitate particle selection in single particle electron microscopy. J Struct Biol 166, 205–213 (2009).

83. S. H. Scheres, RELION: implementation of a Bayesian approach to cryo-EM structure determination. J Struct Biol 180, 519–530 (2012).

84. D. C. Ekiert et al., A highly conserved neutralizing epitope on group 2 influenza A viruses. Science 333, 843–850 (2011).

85. M. Yuan et al., A highly conserved cryptic epitope in the receptor binding domains of SARS-CoV-2 and SARS-CoV. Science 368, 630–633 (2020).

86. Z. Otwinowski, W. Minor, Processing of X-ray diffraction data collected in oscillation mode. Methods Enzymol 276, 307–326 (1997).

87. W. Kabsch, Integration, scaling, space-group assignment and post-refinement. Acta Crystallogr D Biol Crystallogr 66, 133–144 (2010).

88. A. J. McCoy, et al., Phaser crystallographic software. J Appl Crystallogr 40, 658–674 (2007).

89. D. Schritt, et al., Repertoire Builder: high-throughput structural modeling of B and T cell receptors. Mol. Sys. Des. & Eng. 4, 761–768 (2019).

90. P. D. Adams et al., PHENIX: a comprehensive Python-based system for macromolecular structure solution. Acta Crystallogr D Biol Crystallogr 66, 213–221 (2010).

91. P. Emsley, B. Lohkamp, W. G. Scott, K. Cowtan, Features and development of Coot. Acta Crystallogr D Biol Crystallogr 66, 486–501 (2010).

92. A. Shlemov et al., Reconstructing Antibody Repertoires from Error-Prone Immunosequencing Reads. J of Immunol. 199, 3369–3380 (2017).

93. M. A. Larkin et al., Clustal W and Clustal X version 2.0. Bioinformatics 23, 2947–2948 (2007).

94. R. M. Moore, A. O. Harrison, S. M. McAllister, S. W. Polson, K. E. Wommack, Iroki: automatic customization and visualization of phylogenetic trees. PeerJ 8, e8584 (2020).

